# Paraoxonase-like APMAP maintains endoplasmic reticulum-associated lipid and lipoprotein homeostasis

**DOI:** 10.1101/2024.01.26.577049

**Authors:** Blessy Paul, Holly Merta, Rupali Ugrankar-Banerjee, Monica Hensley, Son Tran, Goncalo Dias do Vale, Jeffrey G McDonald, Steven A. Farber, W Mike Henne

## Abstract

Oxidative stress perturbs lipid homeostasis and contributes to metabolic diseases. Though ignored compared to mitochondrial oxidation, the endoplasmic reticulum (ER) generates reactive oxygen species requiring antioxidant quality control. Using multi-organismal profiling featuring *Drosophila*, zebrafish, and mammalian cells, here we characterize the paraoxonase-like APMAP as an ER-localized protein that promotes redox and lipid homeostasis and lipoprotein maturation. APMAP-depleted mammalian cells exhibit defective ER morphology, elevated ER and oxidative stress, lipid droplet accumulation, and perturbed ApoB-lipoprotein homeostasis. Critically, APMAP loss is rescued with chemical antioxidant NAC. Organismal APMAP depletion in *Drosophila* perturbs fat and lipoprotein homeostasis, and zebrafish display increased vascular ApoB-containing lipoproteins, particles that are atherogenic in mammals. Lipidomics reveals altered polyunsaturated phospholipids and increased ceramides upon APMAP loss, which perturbs ApoB-lipoprotein maturation. These ApoB-associated defects are rescued by inhibiting ceramide synthesis. Collectively, we propose APMAP is an ER-localized antioxidant that promotes lipid and lipoprotein homeostasis.

**Key findings summary:**

- APMAP localizes primarily to the ER network in human cells and *Drosophila* fat body tissue, and is a type II integral membrane protein
- Loss of APMAP or *Drosophila* APMAP (dAPMAP) causes ER membrane expansion, elevates CHOP-associated ER stress, promotes LD accumulation, and alters ApoB-lipoprotein secretion
- APMAP-depleted cells and dAPMAP-depleted *Drosophila* fat tissue exhibit defective redox homeostasis; phenotypes associated with APMAP loss are rescued by antioxidant NAC
- Zebrafish-based LipoGlo reporter reveals that loss of *apmap* in zebrafish causes increased vascular ApoB-containing lipoproteins
- Lipidomic profiling indicates that APMAP loss reduces PUFA-phospholipids and elevates intracellular ceramides, which perturbs ApoB maturation

## Introduction

Cellular oxidative stress perturbs organelle and lipid homeostasis, and contributes to pathologies in neurodegenerative diseases, cancers, and metabolic diseases such as non-alcoholic fatty liver disease (NAFLD).^1,2^ To suppress and regulate redox stress, cells express antioxidants including paraoxonase (PON) family enzymes. Humans encode three PON enzymes, of which PON1 is the most well characterized. PON1 is expressed and secreted by liver cells and associates with circulating lipoproteins, where it is thought to serve as an antioxidant esterase by hydrolyzing lipid peroxides bound to lipoproteins HDL and LDL.^3^ Consistent with this, PON1-deficient mice exhibit signatures of atherosclerosis including elevated LDL oxidation, foam cell accumulation, and increased aortic plaques.^4^ PON2 is also highly expressed in liver, and its loss contributes to steatosis.^5,6^ PON3 is primarily expressed in liver and acts as an antioxidant and anti-inflammatory that protects HDL particles from oxidation.^7^

While PON enzymes primarily function in (or are secreted by) liver cells, the PON-like protein C20orf3/APMAP (Adipocyte Plasma Membrane Associated Protein) is expressed in many tissue types. APMAP was initially studied in 3T3-L1 adipocytes, where its siRNA silencing perturbed adipocyte differentiation.^8^ Similarly, APMAP knockout mice are viable but exhibit reduced fat storage and small adipocytes in a diet-induced obesity model.^9^ Despite its early links to adipose tissue, transcript analysis indicates APMAP is ubiquitously expressed, with highest expression in secretory tissues like the liver and thyroid^9^, (also see also ProteinAtlas.org). Furthermore, APMAP over-expression has been noted in cancer cells and correlates with increased hepatic metastasis.^10^ Its elevated expression is linked with increased tumor survival and metastasis, suggesting pro-survival benefits through unclear mechanisms.^11,12^ Structurally, APMAP is a predicted integral membrane protein with a C-terminal six-bladed beta-propeller region similar to the PON1 arylesterase domain. In line with this, human APMAP has been reported to exhibit arylesterase activity *in vitro*, but its *in vivo* function remains unknown.^13,14^ It has been postulated to act as a cellular antioxidant, but this is not experimentally tested.^13^ Furthermore, APMAP’s role in cell homeostasis, and even its sub-cellular localization remain unclear. Though initially identified and named based on its presence in a membrane fraction believed to be the plasma membrane^8^, more recent reports suggest endoplasmic reticulum (ER) localization.^14^

Here, we dissect the sub-cellular localization and cellular functions of mammalian, *Drosophila,* and zebrafish C20orf3/APMAP, revealing them as conserved ER-localized integral membrane proteins that promote intracellular redox and lipid homeostasis in hepatic cells and tissues. APMAP-depleted mammalian cells exhibit defects in ER morphology and altered neutral and phospholipid metabolism, as well as defects in ApoB lipoprotein secretion and lipid droplet (LD) homeostasis. Similarly, *Drosophila* fat bodies (FBs), the hepatic-adipose hybrid tissue of insects, exhibit altered redox and lipid homeostasis when depleted of *Drosophila* APMAP (dAPMAP). Utilizing an in vivo lipoprotein functional assay (LipoGlo)^15^, we show that APMAP loss in zebrafish leads to increased vascular ApoB-lipoproteins, indicating defective lipoprotein homeostasis. LC-MS/MS lipidomic profiling indicates that APMAP-depleted cells manifest changes in phospholipid saturation and ceramide metabolism. Specifically, APMAP loss correlates with increased phospholipid saturation, suggesting lipid remodeling that reduces PUFA-containing lipids, which are preferentially oxidized. We also find APMAP-depleted cells exhibit elevated ceramide levels (commonly increased during oxidative stress), which tightly correlates with defects in ApoB secretion. Critically, defects observed in APMAP-depleted cells can be rescued by treating cells with the chemical anti-oxidant N-acetyl-cysteine (NAC), or by reducing ceramide production, suggesting APMAP loss perturbs cellular redox homeostasis. Collectively, we propose that APMAP is an ER-localized antioxidant that promotes lipid and redox homeostasis to support ER function.

## Results

### APMAP is an ER-localized protein in human cells and *Drosophila* fat body tissue

Previous work indicates APMAP is an integral membrane protein with a predicted N-terminal transmembrane (TM) region and C-terminal arylesterase-like beta-propeller domain (**Fig 1A, B**). In support of this, Alphafold2 predictions indicate the APMAP arylesterase-like domain folds very similarly to the paraoxonase PON1 (**Fig 1C**). Additionally, sequence alignment APMAP and PON enzymes confirm their similarities and a conservation of catalytic residues (**SFig 1A**). Whereas PON1 is soluble and secreted from hepatocytes, APMAP is proposed to remain in cells and co-purify in a cell membrane fraction^8^, but its sub-cellular localization is unclear. To better understand where APMAP localizes in cells, we generated an endogenously mNeonGreen (mNG)- tagged APMAP U2OS homozygous clonal cell line using CRISPR-Cas9 knock-in approaches (**SFig 1B**). This APMAP-mNG fusion decorated the ER network and nuclear envelope, colocalizing with an ER network marker (**Fig 1D**). This is consistent with previous recent studies reporting APMAP localization at the ER network.^14^ This is also consistent with recent organelle-based proteome analysis indicating endogenous APMAP was detected in the ER proteome (https://organelles.czbiohub.org/).

**Figure 1.**
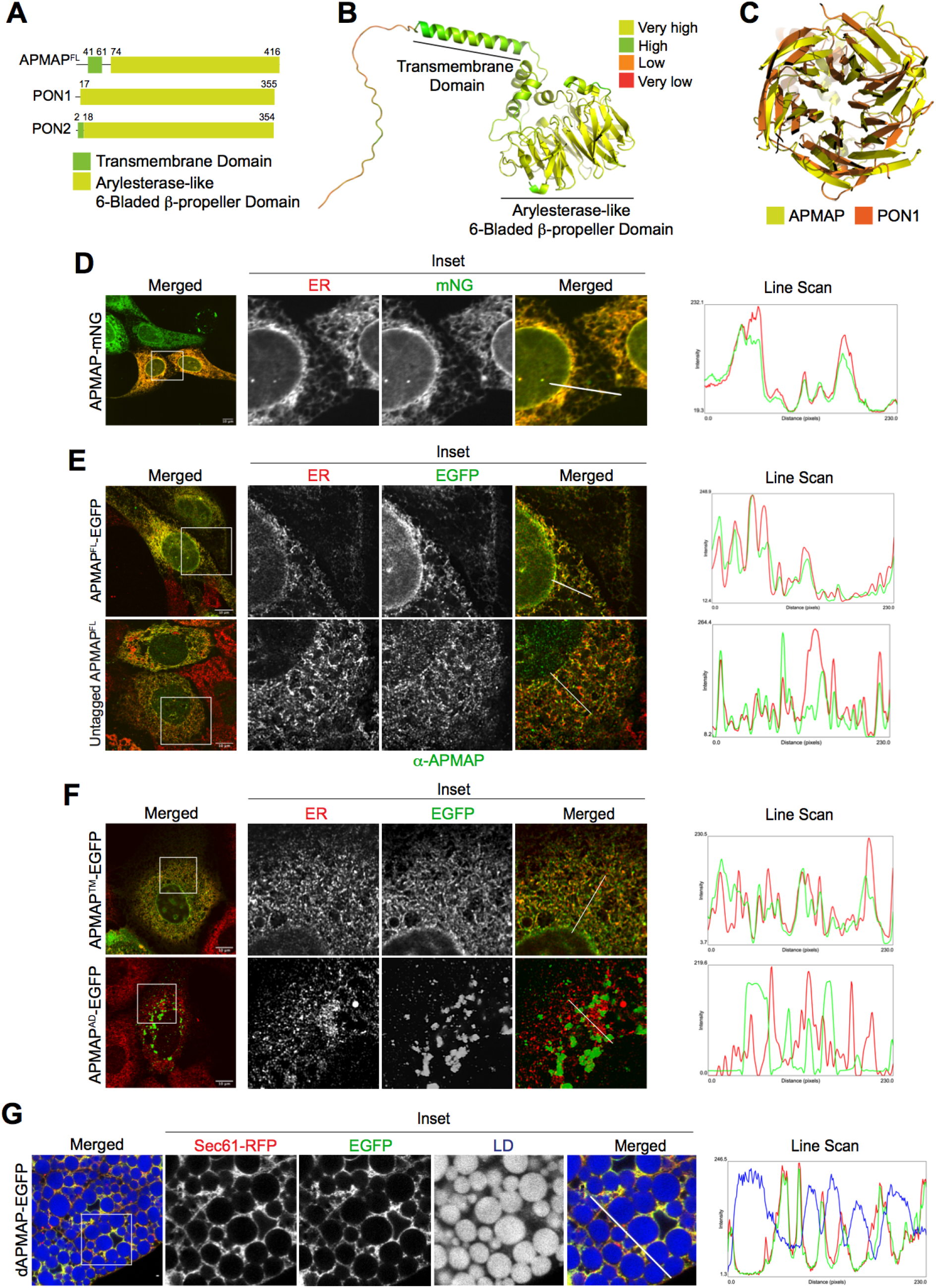
APMAP localizes to the ER in mammalian cells and *Drosophila* fat body. A. Schematic diagram of APMAP domains. APMAP^FL^ depicts the full-length human APMAP consisting of N-terminal transmembrane domain (TM) and C-terminal 6-bladed β-propeller region. PON1 and PON2 are full-length Paraoxonase enzymes with 6-bladed β-propeller domain B. Tertiary structural prediction of human APMAP using AlphaFold (Q9HDC9), highlighting the TM and 6-bladed β-propeller region. The prediction is colored-coded based on model confidence score and assigned color is labelled on the right C. Superposition of the 6-bladed β-propeller regions of AlphaFold predicted APMAP (Q9HDC9) and experimental crystal structure of PON1 (PDB code:1v04), generated using PyMOL (The PyMOL Molecular Graphics System, Version 2.5.5 Schrodinger, LLC) D. Live cell confocal micrographs of C-terminally mNeonGreen (mNG) tagged endogenous APMAP in U2OS cell line showing ER-localization. Cells were transfected with DsRed-KDEL to mark ER network. Scale bar = 10 μm E. Airyscan confocal micrographs of Huh7 cells overexpressed with APMAP-EGFP (Top) and untagged APMAP (Bottom), IF stained using α-HSP90B1 antibody (ER marker, red). Untagged APMAP was co-IF stained using α-APMAP antibody (green) and α-HSP90B1 antibody (ER marker, red). Scale bar = 10 μm F. Airyscan confocal micrographs of Huh7 cells transfected with APMAP^TM^-EGFP and APMAP^AD^-EGFP IF stained with α-HSP90B1 (ER marker, red). Scale bar = 10 μm G. Confocal micrographs of EGFP-tagged *Drosophila* APMAP (dAPMAP) expressed specifically in fat body (FB) using *Dcg-Gal4*>*UAS-*dAPMAP-EGFP. Tissue was co-transfected with ER marker TdTomato-Sec61 (Red) and LDs were stained with MDH (blue). Scale Bar = 10 μm Line scans represent spatial distribution of APMAP (green) with respect to ER maker (red) and LDs (blue). Straight line of 5 pixel width were drawn at ER tubular network and quantified using ‘plot profile’ function in ImageJ.

Since APMAP is reported to be ubiquitously expressed, but with high expression in the liver^16^, we also examined its localization in Huh7 hepatocarcinoma cells, which exhibit liver cell signatures but are also highly amenable to fluorescence imaging. Plasmid-based expression of an EGFP-tagged APMAP in Huh7 cells localized to the ER network and nuclear envelope (**Fig 1E, top**). Similarly, expression of an untagged APMAP was detected by immunofluorescence (IF) using an anti-APMAP antibody, which displayed ER and nuclear envelope localization, indicating that the C-terminal EGFP tag did not affect subcellular localization (**Fig 1E, bottom**). To confirm that the N-terminal TM region was necessary and sufficient for this ER network targeting, we expressed EGFP-tagged APMAP fragments encoding either the TM region alone (APMAP^TM^) or the C-terminal arylesterase-like domain (APMAP^AD^). As expected, the APMAP^TM^-EGFP construct localized to the ER network, whereas APMAP^AD^-EGFP localized to foci in the cytoplasm, but these generally did not colocalize with the ER, and appeared to be protein aggregates (**Fig 1F**). Collectively, this suggests that human APMAP localizes to the ER network via its N-terminal TM region in multiple human cell types.

APMAP is conserved in *Drosophila melanogaster* as CG3373/Hemomucin (Hmu), a poorly characterized protein associated with *Drosophila* lipoprotein homeostasis with a predicted arylesterase function.^17^ For simplicity, we refer to CG3373 as *Drosophila* APMAP (dAPMAP) here. Since the *Drosophila* gene expression databases FlyAtlas.org and Flybase.org indicate that dAPMAP is broadly expressed but highly expressed in the *Drosophila* fat body (FB), we generated an EGFP-tagged UAS/Gal4 dAPMAP *Drosophila* line (*UAS-dAPMAP-EGFP*), and used the FB tissue-specific driver *Dcg-Gal4* to express dAPMAP-EGFP specifically in the FB, the adipose-liver hybrid tissue in insects.^18^ Similar to mammalian APMAP, dAPMAP-EGFP localized to the ER network in larval FB cells, and co-localized with ER marker Sec61-RFP (**Fig 1G**). Collectively, this indicates that both human and *Drosophila* APMAP localize primarily to the ER network of hepatic cells and tissues.

### APMAP is a Type II integral membrane protein with its C-terminus facing the ER lumen

Next we investigated the topology of APMAP at the ER network. Consistent with its ER localization, APMAP encodes a N-terminal signal peptide sequence, and is predicted to have a type II membrane topology with its C-terminus facing the ER lumen.^8^ In support of this topology, previous work detected N-linked glycosylation sites on residues in the C-terminal region, suggesting APMAP’s C-terminus topologically orients into the ER lumen.^16^ To experimentally determine APMAP membrane topology in the ER network, we conducted a selective detergent solubilization and immuno-labeling assay using U2OS cells expressing APMAP^FL^-EGFP (C-terminally EGFP tagged). First, we treated cells with either 0.05% digitonin, which permeates the plasma membrane (PM) but not ER membrane, or 0.1% Triton X-100, which permeates both the PM and ER (**Fig 2A**). As a control, we expressed an ER lumen-localized mCherry construct (mCherry-KDEL), and immuno-stained for the mCherry using an Alexa488 (green) anti-mCherry antibody. As expected, digitonin treatment did not permit immuno-fluorescence (IF) staining of the mCherry-KDEL in the ER lumen, whereas Triton X-100 enabled the anti-mCherry antibody to label mCherry-KDEL in the ER (**Fig 2B**). Similarly, digitonin-treated cells expressing APMAP-EGFP did not exhibit an anti-EGFP staining in the ER tubular network, whereas Triton X-100 treated cells exhibited robust anti-EGFP ER staining along the ER network. This suggests that, as predicted, the C-terminal EGFP-tagged region of APMAP topologically orients into the ER lumen and is accessible only when cells are Triton X-100 treated (**Fig 2B**). However, it should be noted that while we did not detect an anti-EGFP signal in the ER tubular network following digitonin-treatment, we did detect anti-EGFP immunostaining specifically along the nuclear envelope of APMAP^FL^-EGFP expressing cells. We speculate that this limited nuclear envelope signal may be because the nuclear envelope is more sensitive to digitonin permeabilization. In support of this, other studies note that the nuclear envelope is more sensitive to digitonin permeabilization compared to the rest of the ER network.^19^ We also cannot rule out that this nuclear signal is due to newly synthesized APMAP^FL^- EGFP that is exposed to the cytoplasm prior to its insertion into the ER lumen.

**Figure 2.**
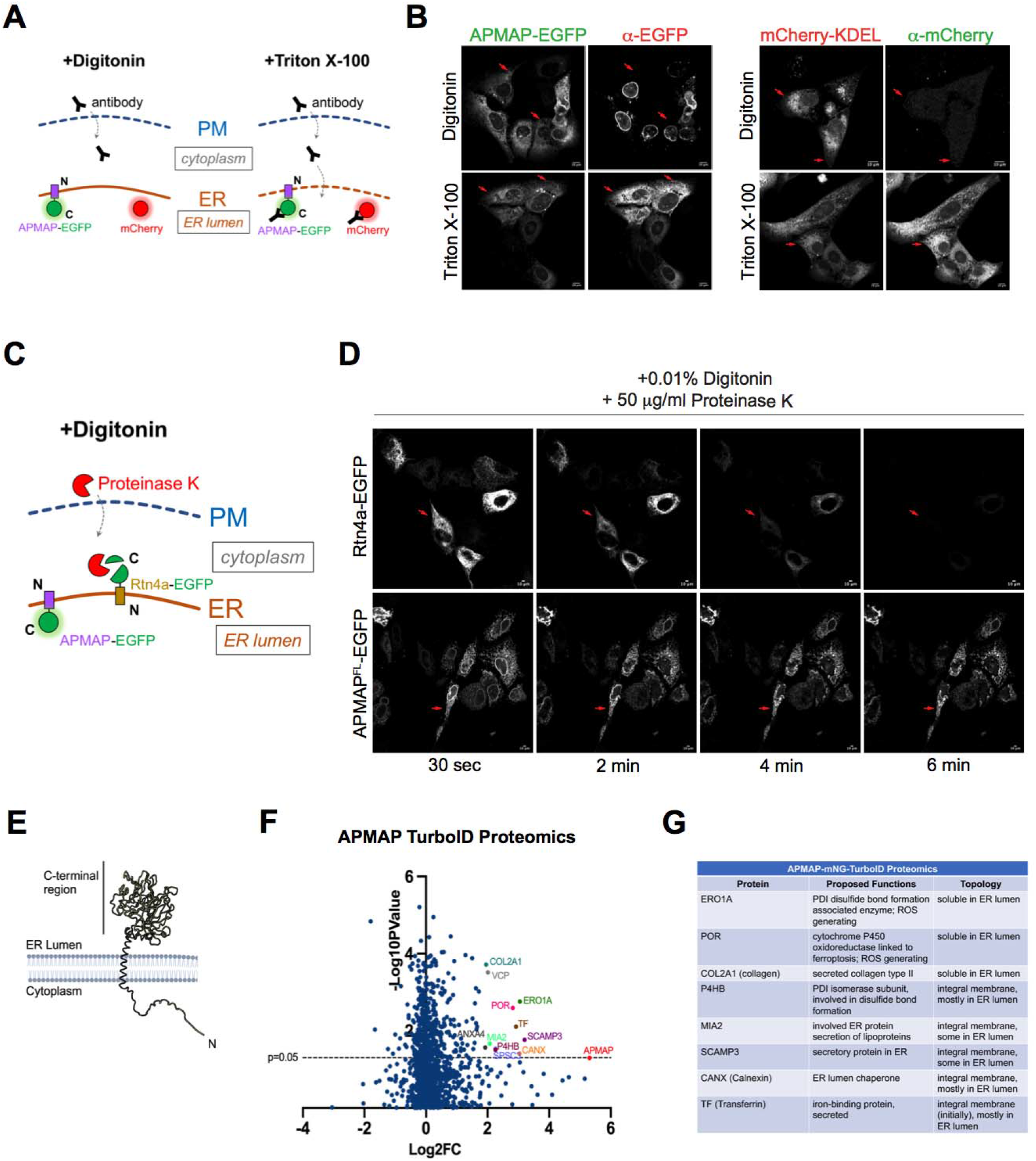
APMAP is a Type II integral membrane protein. A. Cartoon demonstration of semi-permeabilization and immunolabeling experiment for validation of APMAP topology. Cells overexpressed with either APMAP^FL^-EGFP or mCherry-KDEL is treated with either digitonin (permeabilizes only PM) or Triton X-100 (permeabilizes both PM and ER) and are IF stained using α-EGFP and α-mCherry respectively B. Confocal micrographs of U2OS cells expressing APMAP-EGFP or mCherry-KDEL. Cells were treated with either 0.05% digitonin or 0.1% Triton X-100 and IF stained with either α-EGFP (red) and α-mCherry (green) to access exposed C-terminal tag. Scale bar = 10 μm C. Cartoon representation of fluorescence protease protection assay. U2OS cells overexpressed with C-terminally GFP tagged APMAP or Reticulon4a are treated with digitonin to permit selective permeabilization of PM, which is followed by treatment with Proteinase K to allow proteolytic digestion of GFP tag exposed to cytoplasm D. Live cell confocal micrographs of U2OS cells expressing APMAP-EGFP or Reticulon4a-EGFP subjected to subsequent treatments with 0.01% digitonin and 50 μg/ml proteinase K. Images shown are taken 30 sec after treatments up to 6 mins. Scale bar = 10 μm E. Cartoon depiction of the type II membrane topology of human-APMAP, showing N-terminal region facing the cytoplasm and the C-terminal region facing the ER lumen F. Volcano plot of TurboID-based proteomics showing enriched proteins in the APMAP overexpressed Huh7 cells compared to control (untransfected) cells, with Log2 of fold enrichment on X-axis relative to -Log10 of p-values on the Y-axis G. List of selected top protein hits from APMAP Turbo-ID based proteomics, detailing the proposed functions and topology of each gene.

To further test APMAP membrane topology, we conducted detergent permeabilization assays followed by limited proteolysis in living cells. Briefly, living Huh7 cells are treated with 0.01% digitonin together with Proteinase K (PK), and imaged by live-cell confocal laser scanning microscopy. If the EGFP-tag of APMAP^FL^-EGFP orients into the cytoplasm, PK will digest the EGFP and quench its fluorescence immediately following digitonin exposure. In contrast, fluorescence will be preserved if the EGFP faces the ER lumen (**Fig 2C**). Using cells expressing soluble EGFP in the cytoplasm, we confirmed that 0.01% digitonin quickly permeabilizes the PM in live-cell imaging (**SFig 2A**). As a control, we also expressed Reticulon4a-EGFP (Rtn4a-EGFP), which has a known topology with a cytoplasm-oriented EGFP. As expected, following digitonin and PK treatment the Rtn4a-EGFP signal was quickly depleted after ∼6 minutes. In contrast, the APMAP-EGFP signal was retained following digitonin and PK treatment (**Fig 2D**). This again indicates that APMAP-EGFP has a type II membrane protein with its C-terminus facing the ER lumen (**Fig 2E**).

To further dissect APMAP topology, we conducted proximity-based biotinylation labeling by expressing an APMAP construct fused to mNG and the proximity-based biotinylation enzyme TurboID (APMAP-mNG-TurboID).^20^ We confirmed the APMAP-mNG-TurboID construct localized to the ER network, and then conducted biotinylation and affinity purification coupled with LC-MS/MS proteomics to reveal proteins in close proximity to APMAP-mNG-TurboID. Consistent with the C-terminus orienting into the ER lumen, the proteomics dataset revealed numerous proteins localized within the ER lumen (**Fig 2F,G**). These include soluble proteins such as the redox oxidoreductase enzyme ERO1A, collagen type II (COL2Al), as well as ER integral membrane proteins with substantial regions facing the ER lumen such as MIA2/TALI (involved in VLDL secretion), POR (cytochrome P450 oxidoreductase), and PDI subunit P4HB (involved in protein disulfide bond formation and redox homeostasis). Several biotinylated proteins, such as POR, CANX, and PDI, are highly abundant in the ER lumen.^21^ However, other highly enriched proteins in the dataset such as the reactive oxygen species (ROS)-generating enzyme ERO1A are only moderately expressed in the ER, suggesting possible functional interaction, which we explore below. Collectively, this supports a model where APMAP is a type II integral membrane protein with its C-terminus exposed to the ER lumen.

### Loss of APMAP perturbs ER morphology and homeostasis

Since PON family enzymes like PON1 have established roles in the lipid metabolism of lipoproteins, we next investigated whether depletion of APMAP altered lipid homeostasis in the ER network. We RNAi-depleted APMAP in Huh7 cells and confirmed efficient siRNA knockdown via quantitative PCR (Q-PCR) and Western blotting (**Fig 3A**). Next, we conducted immunofluorescence imaging of the ER in either control or APMAP-depleted Huh7 cells. Since the ER is composed of both tubules and sheets, we immuno-labeled both using a general ER stain (anti-Calnexin), as well as labeled ER sheets specifically by staining for CLIMP-63.^22^ Notably, siRNA depletion of APMAP led to visually perturbed ER morphology. First, the ER tubular network appeared less uniform and now contained patches of dense sheet-like signal (**Fig 3B, left**). Second, the CLIMP-63 labeled ER sheets appeared disorganized and less confined to the peri-nuclear region (**Fig 3B, right**). In fact, CLIMP-63 immunosignal was more evident in the cell periphery of APMAP-depleted cells, indicating changes to ER morphology. In line with this, co-staining the same cells for both Calnexin and CLIMP-63 revealed more signal overlap between them when APMAP was depleted (**Fig 3C**). Line-scanning revealed more over-lap of the two ER markers in the cell periphery, suggesting ER membrane expansion in APMAP-depleted cells, a signature of ER stress. In line with this, the ratio of CLIMP-63 positive pixels to Calnexin-positive pixels per cell area was significantly higher in APMAP-depleted cells (**Fig 3D**). Since ER membrane expansion has been observed as a cellular response to alleviate ER stress^23^, we also determined whether APMAP loss elevated known ER stress markers. Indeed, APMAP-depleted cells displayed significantly elevated CHOP and spliced XBP1 (XBP1sp) mRNA transcripts, consistent with ER stress (**Fig 3E**).

**Figure 3.**
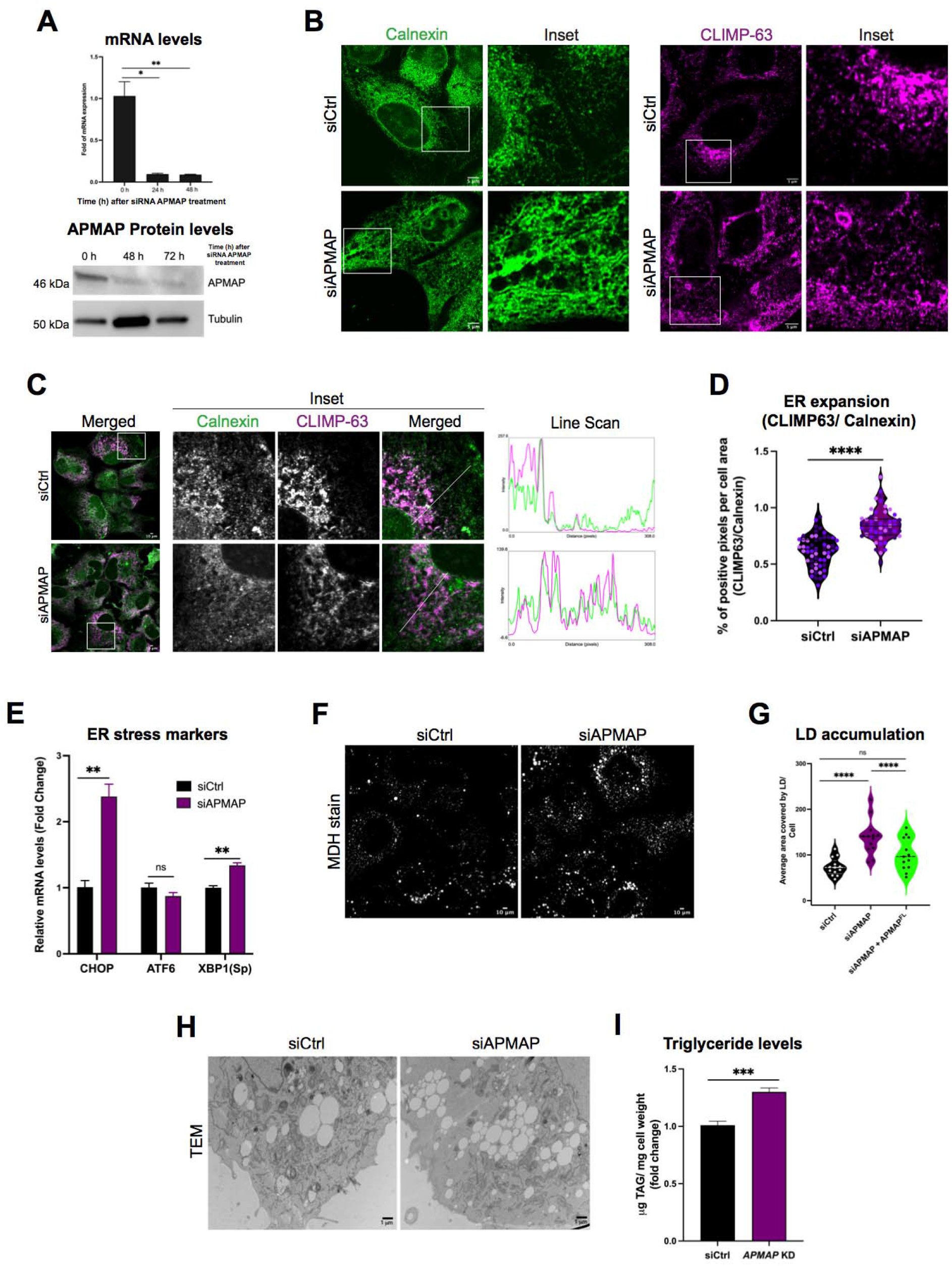
APMAP depletion causes perturbed ER morphology and lipid droplet accumulation. A. (Top) qPCR of APMAP mRNA levels and (Bottom) western blot of APMAP protein levels validating efficient siRNA-mediated APMAP (siAPMAP) knockdown (N=3; **P<0.0021; *P<0.0322; Two tailed unpaired t-test) B. Airyscan confocal micrographs of Huh7 cells treated with scrambled (siCtrl) and siAPMAP. Cells were IF stained using α-Calnexin (green) or α-CLIMP63 (magenta). Scale Bar = 5 μm C. Confocal micrographs of siCtrl and siAPMAP treated Huh7 cells, co-IF stained with α-Calnexin (green) and α-CLIMP63 (magenta). Scale bar = 10 μm. Line scans (straight line with 5 pixel width) representing spatial distribution of calnexin (green) and CLIMP63 (magenta) were produced using “plot profile’ function in ImageJ D. Violin scatter dot plots demonstrating ER membrane expansion in siCtrl and siAPMAP treated Huh7 cells. The plot depicts the ratio of the percent area of CLIMP-63 positive pixels of CLIMP63 over percent positive pixels of Calnexin per cell. Individual data points from more than 50 cells from three sets of independent experiments (**** P<0.0001; Ordinary one-way ANOVA) E. qPCR quantification of mRNA levels of ER stress marker genes (CHOP, ATF6 and spXBP1) from siCtrl and siAPMAP treated cells. N=3, **P<0.0021; Multiple unpaired t-tests Holm-Šídák method with alpha = 0.05 F. Confocal micrographs showing LDs in siCtrl and siAPMAP treated Huh7 cells. LDs were visualized by MDH (gray). Scale bar= 10 μm G. Violin plot representing quantified average area covered by LDs per cell. Total LD area was derived from more than 10 fields of view, each consisting of 10-15 cells and from three sets of experiments (****P<0.0001; one-way ANOVA with Sidak’s multiple comparisons; α = 0.05) H. TEM micrographs of siCtrl and siAPMAP treated U2OS cells followed by a 16h OA treatment to visualize LD morphology. Scale bar= 1 μm I. Quantification of TAG levels in siCtrl and siAPMAP treated cells, using TLC (in micrograms). Data represent mean ± SEM normalized to cell pellet weight; N = 3; ***P<0.0002; Two tailed unpaired t-test

A second indicator of ER stress can be the formation of lipid droplets (LDs) that emerge from the ER network.^24,25^ We stained for LDs using the vital dye monodansylpentane (MDH) and found APMAP-depleted cells exhibited accumulated LDs compared to controls (**Fig 3F,G**). Importantly, this was rescued by re-introduction of an siRNA-resistant APMAP^FL^-EGFP plasmid, showing this effect was due to APMAP loss, as well as demonstrating that EGFP-tagged APMAP retained function (**Fig 3G**).

To determine whether APMAP loss caused LD accumulation in a different cell type, we also depleted APMAP in mouse AML12 hepatocytes. We either transcriptionally silenced APMAP expression using a CRISPR-based plasmid system expressing dCas9 fused with Dnmt3 and KRAB, with an additional cassette gRNA targeting the APMAP promoter (APMAP^KRAB^), or with CRISPR-Cas9-based gene KO (APMAP^KO^). Western blotting confirmed that both strategies led to APMAP depletion in AML12 cells (**SFig 3A**). Similar to Huh7 cells, APMAP depletion in AML12 hepatocytes led to significant LD accumulation (**SFig 3B,C**). We also conducted transmission electron microscopy (TEM) of control and APMAP depleted U2OS cells, which indicated clear LD accumulation when APMAP was lost (**Fig 3H**). Consistent with increased LD staining, APMAP-depleted cells exhibited elevated triglyceride (TG) levels, consistent with LD accumulation (**Fig 3I**). Collectively, this indicates that loss of APMAP alters ER morphology and promotes ER membrane expansion, elevates ER stress markers, and promotes LD and TG accumulation.

### dAPMAP-depleted *Drosophila* display elevated TG stores

We next examined whether loss of dAPMAP in *Drosophila* also led to similar alternations in lipid metabolism. Since the FB serves as both the adipose and hepatic tissue of *Drosophila*, we RNAi-depleted dAPMAP specifically in the FB using the tissue-specific *Cg-Gal4* driver and examined LDs. Similar to APMAP loss in mammalian cells, dAPMAP-depleted third instar larval FBs exhibited larger and more densely packed LDs as well as increased TG stores, indicating dAPMAP loss in the FB led to TG accumulation (**Fig 4 A,B**). Female adult flies with FB-specific dAPMAP depletion also displayed elevated TG stores, whereas dAPMAP-depleted male flies displayed only a negligible change in TG (**Fig 4C**). We also RNAi depleted dAPMAP in the whole-animal using the ubiquitous *Da-Gal4* driver. Similar to dAPMAP loss in the FB, global dAPMAP loss led to significant increases in both larval and adult fly TG (**Fig 4D,E**). To determine whether whole-animal loss of dAPMAP impacted organs other than the FB, we also examined the larval gut. Indeed, there was significant TG accumulation in larval guts of whole-animal dAPMAP-depleted larvae (**Fig 4F,G**). Collectively, this suggests that FB-specific or whole-animal depletion of dAPMAP can alter *Drosophila* neutral lipid metabolism, and generally leads to increased TG storage.

**Figure 4.**
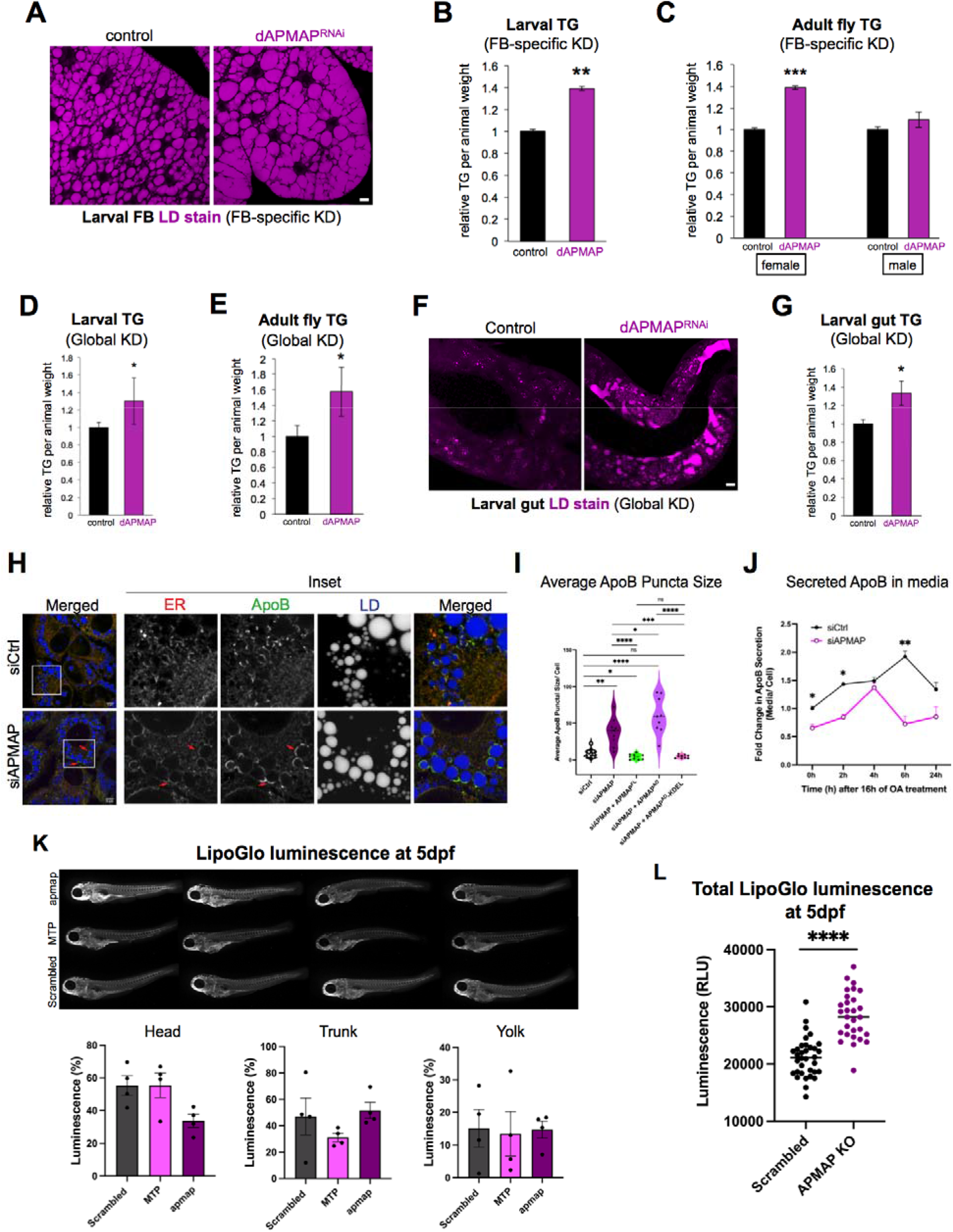
*Drosophila* and zebrafish depletion of APMAP alters TG storage and lipoproteins. A. Confocal micrographs of larval fat-body (FB) tissue sections from control (*Dcg-Gal4* line) and dAPMAP-RNAi line with FB-specific RNAi depletion. LDs are stained with MDH (Magenta). Scale bar = 10 μm B. TLC quantification of relative whole-larval triglyceride (TG) levels from control (*Dcg-Gal4*) and dAPMAP RNAi line with FB-specific RNAi depletion. N=3; **P<0.0021; Two tailed unpaired t-test C. TLC quantification of relative whole adult female and male fly triglyceride (TG) levels from control (*Dcg-Gal4*) and dAPMAP RNAi line with FB-specific RNAi depletion. N=3; ***P<0.0002; Two tailed unpaired t-test D. TLC quantification of relative whole-larval TG levels control (*Da-Gal4*) and dAPMAP RNAi line with global depletion. N=3; *P<0.0322; Two tailed unpaired t-test E. TLC quantification of whole adult fly TG levels from control (*Da-Gal4*) and dAPMAP RNAi line with global depletion. N=3; *P<0.0322; Two tailed unpaired t-test F. Confocal micrographs of larval gut tissue sections from control (*Da-Gal4* line) and dAPMAP-RNAi line with global depletion. LDs are stained with MDH (Magenta). Scale bar = 10 μm G. TLC quantification of relative whole-larval gut triglyceride (TG) levels from control (*Da-Gal4*) and dAPMAP RNAi line with global depletion. N=3; *P<0.0322; Two tailed unpaired t-test H. Airyscan confocal micrographs of siCtrl and siAPMAP treated Huh7 cells, followed by 16 h of OA treatment. Cells were co-IF stained with anti-HSP90B1 (ER marker, Red) and α-ApoB (Green). LDs were stained with MDH (Blue). Red arrows highlight the ApoB crescents situated around LDs. Scale Bar = 10 μm I. ApoB rescue experiments in APMAP depleted cells by re-addition of APMAP^FL^, APMAP^AD^ (cytoplasmic) and APMAP^AD^-KDEL (ER-lumen targeted), all tagged with EGFP. Violin scatter dot plot represents average area covered by ApoB per cell (n >40 cells; N=3; ****P<0.0001, ***P<0.0002, **P<0.0021, *P<0.0332; One-way ANOVA with Sidak’s multiple comparisons; α = 0.05) J. Fold change of ApoB secreted in the culture medium from siCtrl and siAPMAP treated Huh7 cells incubated with OA, determined by ELISA assay. K. LipoGlo luminescence micrographs of 5 days post fertilization (5dpf) showing distribution of ApoB-containing lipoproteins (LipoGlo) in larval zebrafish vasculature. Animals were either treated with non-targeting CRISPR-based guides (scrambled) or gRNAs directed to delete zebrafish APMAP or MTP. Luminescence quantifications for head, trunk, and yolk displayed from N=4 independent experiments. L. Total LipoGlo signal from 5dpf whole-animal lysates. N=3, ****P<0.0001.

We next examined if dAPMAP over-expression altered *Drosophila* lipid homeostasis. While we did not detect significant changes in TG when we over-expressed dAPMAP specifically in the FB, we noted that whole-animal dAPMAP over-expression led to significantly reduced whole-body TG in larvae (**SFig 4A**). Of note, this was the opposite effect of dAPMAP depletion. Similarly, we also observed significant TG reduction in adult female flies globally over-expressing dAPMAP on the *Da-Gal4* driver (**SFig 4B, C).** Male adult flies also demonstrated slightly reduced TG levels, but this trended just below statistical significance. Collectively, we conclude that dAPMAP depletion or over-expression can alter neutral lipid homeostasis, with dAPMAP loss generally associated with TG accumulation similar to APMAP depletion in mammalian cells.

### *Drosophila* tissue and mammalian APMAP depletion alters lipoprotein homeostasis

Since PON family enzymes can associate with circulating lipoproteins, we next examined whether APMAP loss impacted lipoprotein homeostasis. We first stained larval FBs for *Drosophila* lipophorin (Lpp), the major insect lipoprotein that is synthesized and secreted from the FB into the blood (hemolymph).^26,27^ Pre-wandering third instar larvae with FB-specific dAPMAP depletion exhibited a mild elevated intracellular staining pattern for Lpp, suggesting some aspect of Lpp homeostasis was altered with dAPMAP depletion (**SFig 4D**). Relatedly, previous work using purified dAPMAP (i.e. Hmu) indicated it could interact biochemically with Lpp particles isolated from *Drosophila* hemolymph, implying functional crosstalk between dAPMAP and Lpp particles.^17^

Next, we examined lipoprotein homeostasis in Huh7 cells by immunostaining for the major lipoprotein ApoB. Similar to the *Drosophila* FB, we observed that APMAP RNAi-depleted human cells displayed accumulated intracellular ApoB punctae and aggregate-like structures compared to controls (**Fig 4H, I**). In particular, we noted that APMAP deficient cells exhibited “crescent” shaped ApoB puncta closely associated with LDs (**Fig 4H**). Similar ApoB “crescents” have been observed in hepatocytes manifesting ER lipotoxic stress and ApoB accumulation^28^, suggesting APMAP loss perturbed some aspect of ApoB homeostasis. Importantly, we confirmed that re-expression of EGFP-tagged full length APMAP (APMAP^FL^) rescued this ApoB accumulation (**Fig 4I**). To confirm whether APMAP loss in other hepatocytes led to ApoB accumulation, we also stained for ApoB in control and APMAP-depleted AML12 cells. Indeed, APMAP loss led to similar ApoB intracellular accumulation (**SFig 4E**). As an additional comparative control, we also perturbed ApoB folding in the ER by RNAi-depleting ERO1A, which is necessary for disulfide bond formation during ApoB protein folding. Indeed, ERO1A loss caused similar intracellular ApoB accumulation as APMAP depletion, suggesting APMAP is necessary for some aspect of ApoB homeostasis in the ER (**SFig 4F**).

Since APMAP’s C-terminal arylesterase-like domain faces the ER lumen, we also queried whether it needed to localize in the ER to rescue the ApoB accumulation in APMAP-depleted cells. Using Huh7 cells, we expressed the soluble APMAP arylesterase-like domain either in the ER lumen by fusing it to a signal sequence and KDEL (APMAP^AD^-KDEL), or localized it to the cytoplasm (APMAP^AD^) and monitored ApoB accumulation in Huh7 cells. Strikingly, only the ER lumen localized APMAP^AD^-KDEL construct rescued ApoB defects following endogenous APMAP loss (**Fig 4I**). This indicates that APMAP’s primary site-of-action is the ER lumen, and loss of ER lumen-oriented APMAP alters some aspect of ApoB homeostasis.

To determine whether APMAP loss impacted ApoB secretion, we monitored extracellular ApoB protein levels in the Huh7 cell media using an established ELISA-based assay. To stimulate Huh7 ApoB lipoprotein secretion, we treated cells with oleic acid (OA) for 16hrs, then removed the OA and monitored ApoB secretion into the media in a time dependent manner.^29,30^ Consistent with their increased intracellular ApoB immuno-signal, APMAP-depleted Huh7 cells exhibited a significantly lower ApoB secretion into the cell media over time (**Fig 4J**). Collectively, this suggests that APMAP loss alters some aspect of ApoB-lipoprotei homeostasis that impacts their maturation and/or secretion.

### Loss of zebrafish *apmap* increases plasma ApoB-containing lipoprotein levels throughout the larval vasculature

To better understand how APMAP contributes to lipoprotein homeostasis and circulation in the blood, we examined how *apmap* loss influenced atherogenic ApoB-containing lipoproteins in zebrafish larvae, which encode an ApoB homolog and also contain a vascularized circulatory system. We took advantage of the LipoGlo reporter system, which features a luminescent luciferase enzyme (NanoLuc) fused to ApoB (ApoB-NanoLuc, denoted as LipoGlo here) that was inserted in the endogenous *apobb.1* gene locus. LipoGlo can be used to monitor ApoB-containing lipoprotein levels and distribution within zebrafish vasculature.^15^ Indeed, chemiluminescent microscopy at 3 days post fertilization (3dpf) showed that *apmap* depleted larvae exhibited elevated luminescent LipoGlo signal throughout the animal (**SFig 4G**). Zebrafish larvae were also examined at 5dpf, again showing elevated LipoGlo signal in *apmap* depleted larvae (**Fig 4K**). In line with this, whole-animal LipoGlo luminescence was significantly increased in *apmap* depleted larvae compared to scrambled controls, consistent with ApoB-lipoprotein accumulation in the animal vasculature system (**Fig 4L**). A similar accumulation in plasma ApoB-containing lipoproteins was previously observed when *apoC2*, a cofactor in lipoprotein lipolysis, was deleted.^15^ Collectively, these data suggest that *apmap* loss in zebrafish larvae perturbs some aspect of lipoprotein homeostasis that leads to increased total numbers of ApoB-containing lipoprotein particles within the vasculature, a potential atherogenic-like signature.

### APMAP loss alters cellular redox homeostasis

Mechanistically, how does APMAP loss perturb cellular and lipoprotein homeostasis? PON1 is proposed to function as an antioxidant on circulating lipoprotein particles, and its loss drives the accumulation of oxidized lipoproteins and atherogenic pathology.^4^ Similarly, PON2 loss in mice results in enlarged atherogenic lesions and increased oxidative stress markers.^31^ Given APMAP’s similarity to PON enzymes and presence in the ER network, we hypothesized APMAP may function as an ER-localized antioxidant to ensure cellular redox and lipid homeostasis, both critical for lipoprotein synthesis and homeostasis. To test this, we investigated whether APMAP-depleted U2OS cells exhibited increased lipid oxidation by staining them with the fluorescent lipid peroxidation probe BODIPY-C11(581/591). BODIPY-C11 emits red channel fluorescence in its reduced form and displays green 510nm-shifted fluorescence emission when oxidized, thus serving as a membrane-associated oxidation biosensor.^32,33^ BODIPY-C11 stained U2OS cell membranes, but we noted a prominent difference in its oxidized green signal at the cell surface (**Fig 5A**). Of note, the cell surface is exposed to cellular oxidants, but is also where membranes derived from the ER are efficiently delivered from the ER through the secretory pathway. Recent work has also revealed that lipid peroxidation can originate in the ER and then accumulate at the PM.^34^ Line-scanning across the PM indicated that BODIPY-C11 was generally more oxidized at the cell surface in APMAP-depleted cells compared to siRNA control cells (**Fig 5B**). This difference was exacerbated when cells were treated with the chemical oxidant tert-butyl-hydrogen peroxide (TBHP). Indeed, quantification revealed that APMAP-depleted cells exhibited significantly elevated oxidized BODIPY-C11 signal at the cell surface compared to control samples, mimicking control cells exposed to TBHP (**Fig 5B,C**). This suggested APMAP loss altered some aspect of redox homeostasis that correlated with increased oxidation of cellular membranes.

**Figure 5.**
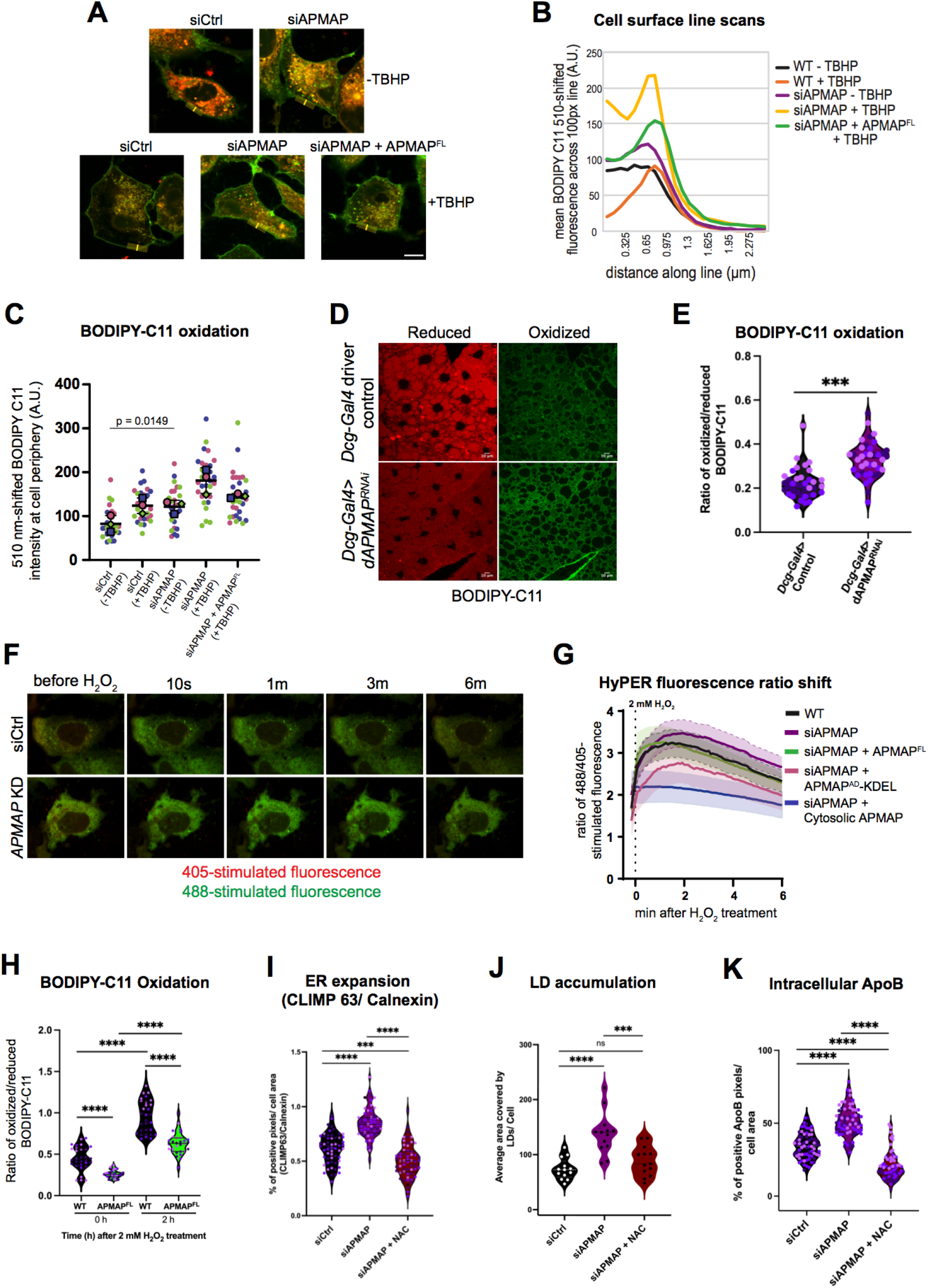
APMAP loss alters cellular redox homeostasis, which is rescued by chemical antioxidant NAC. A. Confocal micrographs of U2OS cells treated with siCtrl, siAPMAP and siAPMAP rescued with siRNA resistant APMAP^FL^ overexpression, incubated with and without TBHP. Cells were stained with BODIPY-C11 to determine the level of oxidation under each condition; Red, reduced form of BODIPY-C11 and Green, oxidized form of BODIPY-C11 B. Line scan (line width of 100 pixels) quantification of 510 nm shift (green; oxidized BODIPY-C11) on the cell surface of siAPMAP treated U2OS cells compared to siCtrl cells C. Fluorescent integrated density of 510 nm shift (green; oxidized) in BODIPY-C11 at the cell periphery was quantified using ImageJ. The statistical significance was assessed by one-way ANOVA with Sidak’s multiple comparison (p=0.0149; 30 cells per condition; N=3). Means +/- SDs shown D. Confocal micrographs of larval FB tissue sections from control (*Dcg-Gal4* line) and dAPMAP-RNAi line with FB-specific RNAi depletion, stained with BODIPY-C11. Images represent oxidized (green) BODIPY-C11 staining and reduced (red) BODIPY-C11 staining in RNAi depleted tissue compared to control tissues. Scale bar= 10 μm E. Quantification of oxidized over reduced BODIPY-C11 in of larval FB tissues represented in Fig 5D was quantified using ImageJ. ***P<0.0021; Two-way ANOVA with Sidak’s multiple comparison _F._ Live confocal micrographs of siCtrl and siAPMAP treated Huh7 cells expressed with ROS biosensor HyPer-ER lumen (denoted HyPER here) construct showing 405 and 488 stimulated fluorescence for 6 mins after treatment with 2mM H_2_O_2_ G. Line graph of fluorescence emission of 488/405 ratio shift measured using HyPer-ERlum in siCtrl and siAPMAP treated Huh7 cells incubated with 2 mM H_2_O_2_. Rescue experiments using different APMAP fragments also shown. Line graphs represent mean of +/- 95% confidence interval (CI) of fluorescence shift. CIs of all conditions shown in dotted lines H. Quantification of oxidized over reduced BODIPY-C11 from WT and APMAP^FL^ overexpressed U2OS cells at 0 and 2 h after 2 mM H_2_O_2_ treatment, were quantified using ImageJ. The statistical significance was assessed by Ordinary one-way ANOVA with Sidak’s multiple comparison (****P<0.0001; n = 30; N=3). Means +/- SEMs shown I. Violin scatter dot plots demonstrating rescue experiment of ER membrane expansion in siAPMAP Huh7 cells by treating with 200uM NAC for 16 h. The plot depicts the ratio of the percent area of CLIMP-63 positive pixels over percent positive pixels of Calnexin per cell. Individual data points from more than 50 cells from three sets of independent experiments (**** P<0.0001; ***P<0.0002; Ordinary one-way ANOVA with Sidak’s multiple comparisons; α = 0.05) J. Rescue experiment of LD morphology in siAPMAP Huh7 cells treated using ROS inhibitor NAC for 16 h. Violin scatter dot plots represent average area covered by LDs per cell in a given ROI. Total LD area was derived from more than 10 ROI, each consisting of 10-15 cells from three sets of experiments (N =3, ****p<0.0001; ***P<0.0002; one-way ANOVA with Sidak’s multiple comparison) K. Violin scatter dot plots demonstrating rescuing of intracellular ApoB levels in siAPMAP Huh7 cells through NAC treatment for 16 h. Each dot depicts the percent area of ApoB positive pixels per cell. (n>50; N=3; **** P<0.0001; Ordinary one-way ANOVA with Sidak’s multiple comparisons; α = 0.05)

Next, we interrogated whether dAPMAP loss impacted *Drosophila* larval FB redox homeostasis using BODIPY-C11. Similar to human U2OS cells, FB-specific dAPMAP depletion in larvae resulted in elevated oxidized (green) BODIPY-C11 staining and reduced red-channel BODIPY-C11 signal compared to control tissues (**Fig 5D,E**). This suggests that dAPMAP loss alters redox homeostasis similar to human APMAP depletion.

Since APMAP localizes to the ER, we also evaluated how its loss impacted redox homeostasis specifically in the ER lumen by expressing an ER lumen-localized version of the ROS biosensor HyPer (HyPER).^35^ As expected, following a hydrogen peroxide pulse to U2OS cells, HyPER exhibited a fluorescence emission 488/405 ratio shift followed by a gradual return to baseline over several minutes (**Fig 5F,G**). We monitored the response of siRNA control and APMAP-depleted cells to this TBHP insult. Notably, the 488/405 ratio shift was visually more pronounced in APMAP-depleted cells, with an elevated peak ratio shift at ∼2min after TBHP addition, and a milder recovery trajectory compared to control cells (**Fig 5F,G**). Re-introduction of siRNA-resistant APMAP returned the 488/405 ratio curve to a WT trajectory (**Fig 5G**). Cells over-expressing APMAP’s arylesterase-like domain either in the ER lumen (APMAP^AD^-KDEL) or cytoplasm (cytosolic APMAP) displayed significantly reduced 488/405 ratio, indicating that APMAP is sufficient to reduce the oxidative environment following hydrogen peroxide exposure (**Fig 5G**). To further test this, we over-expressed full length APMAP and monitored BODIPY-C11 oxidation in U2OS cells. Indeed, APMAP^FL^ over-expression was sufficient to lower the baseline BODIPY-C11 oxidation in cells, and it significantly reduced BODIPY-C11 oxidation when cells were treated with 2mM hydrogen peroxide for 2 hours (**Fig 5H**). Collectively, this indicates that APMAP loss or over-expression can influence the cellular redox state.

### APMAP loss can be rescued by treatment with chemical antioxidant NAC

We reasoned that APMAP serves as an ER-localized antioxidant. In support of this, the TurboID-based APMAP interactome detected major ER ROS generating enzymes like ERO1A in the local APMAP interactome, suggesting APMAP may be situated in close physical proximity to ER machinery that generates ROS (**Fig 2F**). To test this, we treated control and APMAP-depleted U2OS cells with the chemical antioxidant N-acetyl cysteine (NAC), and determined whether this could rescue defects observed with APMAP loss. First, we monitored whether NAC treatment could suppress the ER membrane expansion observed in Huh7 cells. Indeed, 24hr of NAC treatment suppressed the ER membrane expansion seen in APMAP-depleted cells (**Fig 5I**). Similarly, NAC treatment also completely rescued LD accumulation in APMAP-depleted Huh7 cells (**Fig 5J**). Finally, we found that NAC treatment rescued the ApoB accumulation in APMAP-depleted cells (**Fig 5K**). Collectively, this suggests that APMAP and dAPMAP loss cause perturbations in ER redox homeostasis that promote oxidation. Remarkably, APMAP loss can be rescued by treating cells with the chemical antioxidant NAC. This supports a model where APMAP functions as an ER-localized antioxidant.

### Lipidomic profiling reveals APMAP loss alters lipid metabolism and elevates ceramides

Oxidative stress can alter the cellular lipid profile and lead to fat accumulation, phospholipid peroxidation, and the production of stress signaling lipids such as ceramides.^36^ To evaluate how APMAP loss impacted cellular lipids, we next conducted lipidomic profiling using liquid chromatography coupled with mass spectrometry (LC-MS/MS). Since we wanted to probe how APMAP loss contributed to redox homeostasis, we treated cells with a sub-lethal dosage of TBHP prior to harvesting for lipidomics.

Lipidomics revealed changes in the cell lipid profile with APMAP depletion. There were significant reductions in the relative abundance of several phospholipids including PE as well as lyso-phospholipids like lyso-PE and lyso-phosphatidylcholine (lyso-PC) (**Fig 6A,B, SFig 6A,B**). While total phosphatidylcholine (PC) remained unchanged with APMAP loss (**Fig 6C**), lipidomics revealed alterations in the PC lipid saturation profile. In particular, there was a decrease in the PC polyunsaturated fatty acid (PUFA) content, with a corresponding increase in mono-unsaturated fatty acid (MUFA) content (**Fig 6D**).

**Figure 6.**
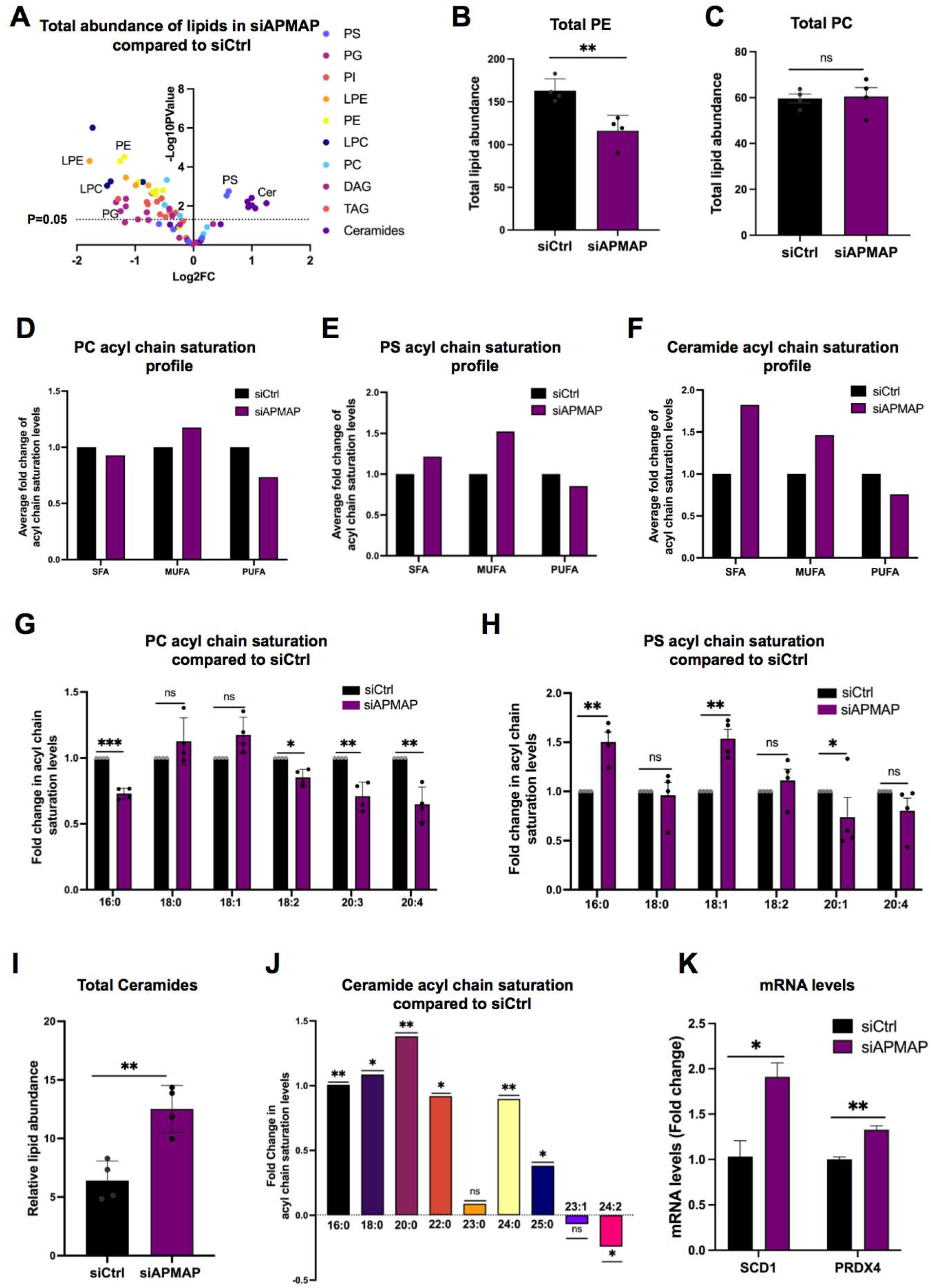
APMAP depletion alters lipid saturation and increases ceramides. A. Volcano plot depicting total abundance of different lipid species in siAPMAP treated Huh7 cells compared to siCtrl cells, determined through LC-MS/MS. Each data point represents the average level of individual fatty acid of a particular lipid species. Log2 fold enrichment of the fatty acid (x-axis) is plotted against the - Log10 transformed p-values from the unpaired t-test (y-axis) B-C.Total lipid abundance of PE and PC respectively in siCtrl and siAPMAP treated Huh7 cells, measured with LC-MS/MS. N=4; **P<0.0021, ns=not significant; two-tailed unpaired t-tests D. Levels of saturated (SFA), monounsaturated (MUFA), and polyunsaturated (PUFA) fatty acyl chains in PC in siCtrl and siAPMAP Huh7 cells represented as average of fold change; N=4 E. Levels of saturated (SFA), monounsaturated (MUFA), and polyunsaturated (PUFA) fatty acyl chains in PS in siCtrl and siAPMAP Huh7 cells represented as average of its fold change; N=4 F. Levels of saturated (SFA), monounsaturated (MUFA), and polyunsaturated (PUFA) fatty acyl chains in ceramides in siCtrl and siAPMAP Huh7 cells represented as average of its fold change; N=4 G. Bar diagram representing the mean fold change (N=4) of saturation index of all acyl chains in PC in siCtrl and siAPMAP treated cells, measured with LC-MS/MS. N=4; ***P<0.0002, **P<0.0021, *P<0.0332; Multiple unpaired t-tests Holm-Šídák method with α = 0.05 H. Bar diagram representing the mean fold change (N=4) of saturation index of all acyl chains in PS in siCtrl and siAPMAP treated cells measured with LC-MS/MS. N=4; ***P<0.0002, **P<0.0021, *P<0.0332; Multiple unpaired t-tests Holm-Šídák method with α = 0.05 I. LC-MS/MS measurement demonstrating total relative lipid abundance of ceramides in siCtrl and APMAP depleted Huh7 cells. N=4, **P<0.0021; Two-tailed Unpaired t-tests J. Quantification of fold change in saturation level of different ceramide fatty acyl chains in siAPMAP treated cells compared to siCtrl (positive values denote higher in siAPMAP), measured with LC-MS/MS. **P<0021, *P<0.0332; Ordinary One-way Anova K. Quantified relative gene expression levels of SCD1 and PRDX4 in siCtrl and siAPMAP treated Huh7 cells, using qPCR. N=3; **P<0.0021, *P<0.0332; Two-tailed Unpaired t-tests

PC species containing 18:2, 20:3, and 20:4 acyl chains were significantly decreased in APMAP-depleted cells (**Fig 6G**). Similarly, phosphatidylserine (PS), another abundant phospholipid, also exhibited a slight decrease in PUFA content and elevated MUFA phospholipids (**Fig 6E**). Indeed, PS 18:1 was significantly elevated, and there was a slight downward trend of PS 20:4 that was just below statistical significance (**Fig 6H**). This collectively indicates that APMAP loss correlates with a decreased PUFA profile for major phospholipids. One possible explanation for this PUFA depletion is the oxidation of PUFA-containing phospholipids when APMAP is depleted (i.e. oxi-PUFA phospholipids would no longer be detected, and observed as reduced PUFA phospholipids). However, we cannot rule out that decreased PUFA synthesis could also contribute to these changes. Since lipid saturation is closely correlated to membrane fluidity and shape, these lipid changes may contribute to the ER morphology alternations observed in APMAP-depleted cells, which displayed sheet-like membrane expansion. Indeed, this may also explain why treatment with the antioxidant NAC rescued the ER morphology defects when APMAP was depleted (**Fig 5I**).

Whereas many phospholipids were unaltered or slightly decreased in total abundance, we noted that APMAP-depleted cells exhibited significant increases in ceramides (**Fig 6A,I**). Ceramides generally contain a fully saturated acyl chain, and conditions that reduce PUFA lipids in favor of MUFA or SFA lipids can result in increased ceramide production.^37^ Elevated ceramide synthesis is also tightly correlated to oxidative stress and ER stress.^38,39^ In line with this, we observed increased PRDX4 transcripts, an antioxidant linked to ER stress response, in APMAP-depleted cells (**Fig 6K**). Lipidomics also revealed that APMAP-depleted cells were enriched in medium and long chain ceramides, including ceramides 16:0, 18:0 and 20:0 that are associated with metabolic dysfunction and oxidative stress (**Fig 6F,J**).^40^

Collectively, lipidomic profiling indicates APMAP-depleted cells exhibit decreased PUFA phospholipids and increased ceramides, both signatures of oxidative stress. PUFA lipid loss may be due to increased PUFA oxidation with APMAP loss. Alternatively, APMAP-depleted cells may also actively remodel their lipid metabolism to reduce synthesis of PUFA-containing phospholipids (which are sensitive to oxidation) in favor of MUFA lipids and ceramides. In support of this model, APMAP-depleted cells exhibit increased SCD1 desaturase mRNA transcripts, which generates MUFA-containing lipids, and is generally elevated in response to lipotoxic stress (**Fig 6K**).

### Intracellular ceramides associate with and perturb ApoB in APMAP-depleted cells

The liver is the primary site of ceramide synthesis, and ceramide accumulation in hepatocytes is a signature of liver stress and inflammation, and can alter lipoprotein secretion and even promote foam cell formation in atherogenic models.^41,42^ Oxidative stress can drive ceramide production as a secondary messenger and stress signal.^43^ Given that APMAP-depleted mammalian cells displayed elevated ceramides by lipidomics, we hypothesized that this may perturb some aspect of ApoB-lipoprotein homeostasis when APMAP was depleted.

To investigate whether ceramides influenced ApoB homeostasis, we directly monitored intracellular ceramides using an established anti-ceramide antibody. As a control, we treated cells with the ceramide synthase (CerS) inhibitor fumonisin B. As predicted, this lowered the anti-ceramide immuno-fluorescence (IF) signal of Huh7 cells (**Fig 7A**). Consistent with lipidomic profiling, ceramide IF staining was higher in APMAP-depleted Huh7 cells compared to control cells (**Fig 7B**). We also observed elevated ceramide IF staining in APMAP-depleted mouse AML12 hepatocytes, indicating this ceramide accumulation was conserved across species and cell types (**SFig 7A**). As expected, we could lower ceramides in APMAP-depleted Huh7 cells by treating them with fumonisin B (F), and even further reduce it through combined treatment with fumonisin B and myriocin (F+M), which inhibits the serine palmitoyltransferase (SPT) complex involved in ceramide production (**Fig 7A,B**).

**Figure 7.**
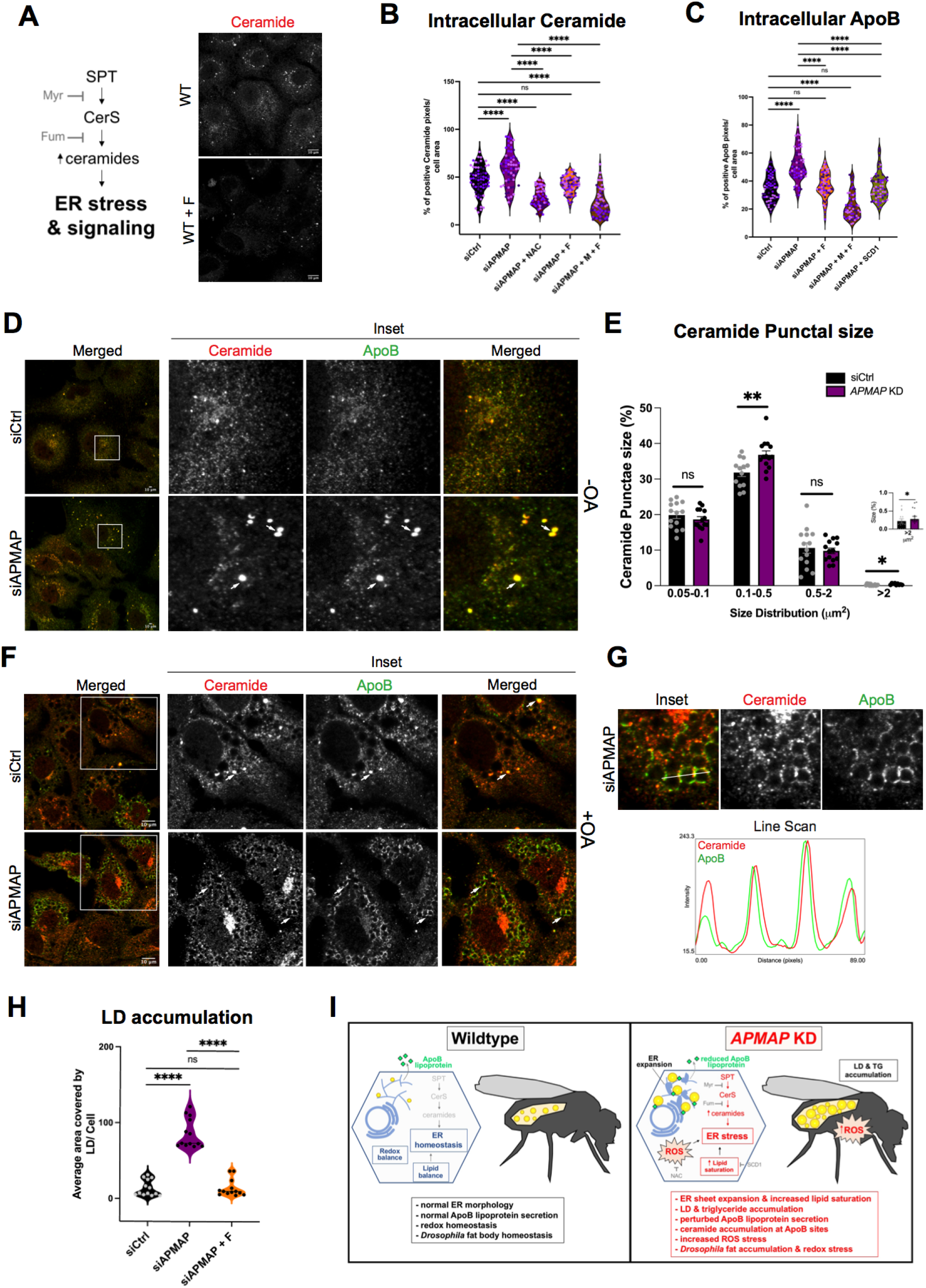
Suppression of ceramide synthesis and NAC treatment in APMAP-depleted cells rescues intracellular ApoB accumulation. A. Schematic representation of ceramide synthesis and its involvement in activation of ER stress and signaling. Myriocin (Myr or M) and Fumonisin B (Fum or F) act as inhibitors of Serine Palmitoyltransferase (SPT) and Ceramide Synthase (CerS) respectively. Representative images of ceramide levels in WT and Fumonisin B treated Huh7 cells B. Violin scatter dot plots of intracellular ceramide levels in Huh7 cells treated with different conditions. Each dot depicts the percent area of ceramide positive pixels per cell. (n=65; N=3; **** P<0.0001; Ordinary one-way ANOVA with Sidak’s multiple comparisons; α = 0.05) C. Violin scatter dot plots demonstrating intracellular ApoB levels in Huh7 cells under different conditions. Each dot depicts the percent area of ceramide positive pixels per cell. (n=65; N=3; **** P<0.0001; Ordinary one-way ANOVA with Sidak’s multiple comparisons; α = 0.05) D. Confocal micrographs of siCtrl and siAPMAP treated Huh7 cells, co-IF stained with α-Ceramide and α-ApoB. White arrows highlight the ceramide puncta co-localized with ApoB. Scale bar = 10 μm E. Histogram quantification of ceramide punctal sizes (% of total punctal population) in a given size range in μm^2^ in siCtrl and siAPMAP treated cells F. Confocal micrographs showing co-localization of ceramide and ApoB in siCtrl and siAPMAP Huh7 cells treated with OA for 16 h. Highlighted areas show the formation of ceramide rings along with ApoB crescents in siAPMAP, compared to punctal-like ceramide structures in siCtrl. Scale bar = 10 μm G. Inset of confocal images from siAPMAP Huh7 cells showing ceramide and ApoB co-localization and their line scan (line width of 10 pixels) demonstrating the spatial distribution H. Violin plots demonstrating the rescue of LD accumulation in siAPMAP cells using Fumonisin. Each dot represents average area covered by LDs per cell in a given ROI. (13 ROI, each consisting of 10-15 cells from three sets of experiments; ****p<0.0001; ***P<0.0002; one-way ANOVA with Sidak’s multiple comparison) I. Working model for role of APMAP in hepatic cell and *Drosophila* tissue ER redox homeostasis, lipid metabolism, and normal ApoB lipoprotein secretion

We next asked whether redox dysfunction in APMAP-depleted cells contributed to ceramide elevation. We treated APMAP-depleted cells with antioxidant NAC. Indeed, this lowered ceramide levels (**Fig 7B**), suggesting the ceramide accumulation in APMAP-depleted cells required defects in redox homeostasis associated with APMAP loss. As a control to determine whether this ceramide elevation was specific to APMAP loss, or would occur with general ER stress, we also RNAi-depleted the ER enzyme ERO1A. Notably, ERO1A loss did not lead to ceramide accumulation like APMAP loss, suggesting ceramide elevation was specific to APMAP depletion (**SFig 7B**).

Ceramides are known to associate with lipoproteins and even traffic on them, and alterations in ceramide metabolism can impact lipoprotein homeostasis and secretion.^44^ Given that APMAP depleted cells manifested defective ApoB secretion, we queried whether ceramide accumulation influenced intracellular ApoB deposits. Strikingly, ApoB accumulation in APMAP-depleted cells was returned to control levels following fumonisin B treatment, and further reduced by dual treatment with fumonisin B + myriocin (**Fig 7C**). Notably, we could also lower ApoB accumulation by over-expressing the fatty acid desaturase SCD1 in siAPMAP cells, consistent with the model that ApoB accumulation was due to altered lipid metabolism when APMAP was lost (**Fig 7C**).

We hypothesized that intracellular ceramide was associating with ApoB deposits. To test this, we co-immunostained for both ceramide and ApoB. Strikingly, the ceramide and ApoB immuno signals tightly overlaid, and we noted ceramide accumulations in enlarged ApoB punctae in APMAP-depleted cells (**Fig 7D, arrows**). Quantification confirmed these ceramide-ApoB punctae were significantly larger in APMAP-depleted cells compared to control cells, suggesting ceramide was directly associated with ApoB accumulations (**Fig 7E**). We also observed these ceramide-ApoB structures when we treated Huh7 cells with OA to stimulate TG synthesis and ApoB-lipoprotein production. Here, ceramide colocalized tightly with the crescent-like ApoB structures near LDs, suggesting ceramide associated with ApoB during periods of ApoB lipidation (**Fig 7F,G inset and linescan**).

Finally, since APMAP loss also led to LD accumulation, we also queried whether lowering ceramides via fumonisin B treatment could rescue this. Indeed, fumonisin B treatment returned LD levels of APMAP-depleted cells to control levels (**Fig 7H**).

Altogether, this indicates that APMAP depleted cells exhibit elevated ceramides that associate with intracellular ApoB and perturb ApoB-lipoprotein homeostasis. It suggests ceramides disrupt some aspect of ApoB lipidation and lipoprotein maturation in the ER, leading to changes in ApoB-lipoprotein homeostasis secretion and lifetime in circulation. This may explain why in mammalian cells there are defects in ApoB release, and in zebrafish there is accumulation of ApoB-containing lipoproteins in circulation. These ApoB-linked defects may also explain the TG and LD accumulation observed in mammalian cells and *Drosophila* fat tissue when APMAP is lost.

### Conclusions and limitations of study

Collectively these experiments support a model where APMAP functions as an ER-localized antioxidant that maintains redox and lipid homeostasis in mammalian cells, *Drosophila,* and zebrafish (**Fig 7I**). APMAP-depleted cells exhibit several signatures consistent with altered redox homeostasis including elevated ER stress, altered ER morphology, defects in ApoB maturation, LD accumulation, and increased cellular oxidation. Depletion of dAPMAP in the *Drosophila* fat body leads to fat accumulation and defects in redox homeostasis. Zebrafish lacking *apmap* manifest increased plasma ApoB-lipoproteins, an atherogenic signature. Mechanistically, we find that APMAP loss alters the cellular lipid profile, including elevating ceramides that associate with ApoB. This appears to be driven by redox stress, as treating APMAP-depleted human cells with antioxidant NAC rescues LD accumulation and ApoB secretion. We propose that with APMAP loss, there is an increase in oxidative stress and lipid oxidation at the ER. Cells respond by elevating ceramides, which in turn alters ApoB homeostasis and secretion from the ER, which also promotes intracellular TG and LD accumulation (**Fig 7I**). A limitation of the study is that we do not know whether ceramide directly inhibits the lipidation of ApoB lipoproteins or some other stage of their maturation or secretion. Ceramides may also act as signaling molecules that alter other aspects of lipoprotein secretion. Future studies will investigate how ceramides influence ApoB and ER homeostasis when APMAP is lost.

## Discussion

Oxidative stress contributes to numerous diseases including NAFLD and cardiovascular diseases.^45,46^ Liver tissue is particularly susceptible to oxidative damage, and as such hepatocytes encode antioxidant defenses that are incompletely understood. Here, we identify APMAP and its *Drosophila* ortholog dAPMAP as antioxidants expressed in diverse cells and tissues, and profile how their loss perturbs lipid and redox metabolism. We find that APMAP loss in *Drosophila* and zebrafish alter lipoprotein homeostasis, and in zebrafish promote increased ApoB-lipoprotein particles in the circulating plasma. We also find that APMAP and dAPMAP are integral membrane proteins that primarily localize to the ER network. In line with this, loss of APMAP in mammalian cells alters ER morphology, resulting in ER membrane expansion, LD and TG accumulation, and perturbed ApoB secretion. Notably, this all these defects are rescued by treating cells with the chemical antioxidant NAC, underscoring how APMAP loss alters cell homeostasis through redox dysfunction. Lipidomic profiling reveals that APMAP-depleted cells have altered lipid saturation profiles featuring reduced PUFA phospholipids and elevated ceramides. Surprisingly, we find that ceramides associate with ApoB in cells and perturb ApoB homeostasis. As such, treating APMAP-depleted cells with ceramide synthesis inhibitors rescues intracellular ApoB accumulation, as well as LD accumulation. Collectively, we propose that APMAP loss causes ER stress that, among other consequences, elevates ceramide pools that perturb some aspect of ApoB maturation.

Defects in lipid metabolism can drive liver diseases including NAFLD and liver steatohepatitis.^47,48^ Liver cells encode antioxidants including glutathione and superoxide dismutases, as well as enzymes that protect lipids from peroxidation such as GPX4.^49^ Generally these factors act in the cytoplasm, mitochondria, or plasma membrane, where lipid peroxidation can fatally damage cellular energetics or membrane integrity. How the ER network is protected against oxidative stress is less understood, though critical to cellular redox homeostasis. Although generally ignored compared to the ROS-generating mitochondria, the ER network can generate ∼25% of all cellular ROS.^50^ This is particularly true in highly secretory cells like hepatocytes or *Drosophila* FB cells. Furthermore, recent studies of lipid peroxidation indicate that the ER is a key site of action for anti-ferroptosis drugs, suggesting ER lipid dysfunction is a driver for ferroptosis.^34^ We propose that APMAP is a previously unappreciated ER-localized antioxidant that contributes to cellular redox and lipid homeostasis.

Mechanistically, how does APMAP loss impact ER homeostasis? We find that APMAP depletion causes accumulation of intracellular ApoB, which is synthesized and folded in the ER lumen prior to its secretion on lipoproteins. These ApoB accumulations co-localize strongly with ceramide, and it is notable that ceramide can associate with ApoB and lipoproteins.^41^ We speculate that the ApoB accumulations in APMAP-depleted cells represent aggregated ApoB protein bound to ceramide that is in the process of being lipidated with TG and phospholipids. We also find by TurboID-based proteomics that ROS producing ER lumen proteins like ERO1A are in close proximity to APMAP.^49^ ERO1A contributes to disulfide bond formation of proteins like ApoB (which contains eight disulfide bonds) during their co-translational folding in the ER.^51^ We speculate that APMAP localizes throughout the ER network, but potentially acts on lipids or other oxidized metabolites in close proximity to ERO1A and ApoB, where local ROS pools are generated during ApoB disulfide bond formation and lipidation. In this model, APMAP acts as a ‘quality control’ factors that ensures lipids or other molecules are not oxidized in the ER during ROS generating processes.

APMAP shares homology with PON family enzymes, which function as antioxidants either on lipoproteins or in cells. In line with this, mice lacking PON enzymes exhibit oxidative stress and atherogenic signatures such as disrupted fat metabolism, foam cell accumulation, and atherosclerosis plaques.^4^ Our observations in animal models supports the hypothesis that APMAP loss promotes similar pathologies. dAPMAP loss in *Drosophila* disrupted lipoproteins and led to fat accumulation in the larval FB, and *apmap* depletion in zebrafish led to increased vascular ApoB-containing lipoproteins, a potential atherogenic-like signature. Future studies will investigate how APMAP contributes to liver homeostasis and lipid/protein quality control in circulation. They will also examine what specific substrate(s) APMAP acts upon. PON enzymes are esterases but display broad substrate specificity, and can act on lactones and organic toxins including paraoxons. It is unclear whether APMAP has similar substrates.

It is notable that APMAP up-regulation has been noted in several cancer models, and its deletion sensitizes cancer cells to phagocytic engulfment.^14^ We speculate that APMAP’s ability to promote ER lipid and redox homeostasis provides cancer cells with a survival advantage when over-expressed. Conversely, APMAP loss may alter the lipid profile of the cancer cell surface, which may promote engulfment by phagocytes, perhaps by increasing the presence of specific oxi-lipids or related molecules on the surface that serve as ‘eat me’ signals for phagocytes. Future studies will no doubt continue to dissect how APMAP influences cellular redox homeostasis and lipid quality in physiology and diseases like cancers.

## Acknowledgements

We thank members of the Henne lab as well as Mike Bassik (Stanford) and Roarke Kamber (UCSF) for helpful advice and discussions during the development of this study. We also especially thank Joan Font-Burgada for assistance in producing the CRISPR-Cas9 constructs for this work. This work is supported by grants from the Welch Foundation I-1873 (W.M.H), the National Institutes of Health (NIH) grants GM119768 (W.M.H), DK126887 (W.M.H.), R01DK093399 (S.A.F.) and R01DK116079 (S.A.F.) the Ara Parseghian Medical Research Fund, and the UT Southwestern Endowed Scholars Program.

## Methods

### Mammalian cell culture

Human cell lines Huh7 and U2OS were cultured in a 37°C incubator with 5% CO_2_ in DMEM high glucose (Sigma D5796) supplemented with 10% heat-inactivated fetal bovine serum (Sigma F4135), 25 mM HEPES (Sigma H0887), and 1% penicillin/streptomycin (Sigma P4333).

The AML12 mouse hepatocyte cells were cultured in DMEM/F-12 50/50 mix with L-glutamine and 15 mM HEPES (Corning 10-092-CV) supplemented with 10% heat-inactivated fetal bovine serum (Sigma F4135), 1x Insulin-Transferrin-Selenium supplement (ITS) (Sigma I3146, 100x), 40 ng/ml dexamethasone and 1% penicillin/streptomycin (Sigma P4333).

0.25% Trypsin-EDTA (Sigma T4049) was used to passage cells when the confluency reaches approximately 90%. Cells were tested for mycoplasma upon the first thawing and upon generation of new cell lines using a standard PCR protocol with oligos (Forward – 5’ CCGCGGTAATACATAGGTCGC 3’; Reverse – 5’ CACCATCTGTCACTCTGTTAACC 3’).

### Chemicals and reagents

For LD biogenesis and cell treatments, the following reagents were used for indicated period of time in each experiment. 1) LD biogenesis with 600 μM of Oleic Acid (OA) conjugated with 100 μM of fatty acid-free BSA (Sigma A3803) for 16-18h, 2) ROS inhibition with 200 μM of N-Acetyl Cysteine (Sigma A9165) for 16-18h, 3) Oxidative stress induction using 70 μM tert-Butyl hydroperoxide (Thermo Fisher 180345000) for 16-18 h, 4) 50 μg/ml Proteinase K (Sigma P2308) and 0.01% or 0.05% digitonin (Sigma 300410) for APMAP topology experiment, and 4) Ceramide rescue experiments using 0.02 mg/ml Fumonisin B1 (Sigma F1147) and 20 μM myriocin (Sigma M1177) for 16-18h.

### Cloning and constructs

Full length APMAP (APMAP^FL^), APMAP^TM(1–77)^ and APMAP^AD(77–416)^ tagged with EGFP at their C-termini were generated after PCR amplification of fragments of interest from full length APMAP, which was amplified from human cDNA library and cloning into pEGFP-N2 using Gibson assembly method (NEB E2611L). A stop codon was inserted before EGFP in pEGFP-N2 vector to clone untagged APMAP^FL^. APMAP^AD^-KDEL (untagged) was made from TagBFP-KDEL (Addgene 49150) by replacing TagBFP between the ER signal sequence of BiP and KDEL ER retention signal with the ER-luminal portion of APMAP. PCR amplified fragments of full-length APMAP, TurboID, and mNeonGreen Gibson Assembling these into the EGFP-N2 backbone without GFP for the construction of APMAP-TurboID-mNeonGreen. The linker between APMAP and TurboID is a 6-glycine linker, and the linker between TurboID and mNeonGreen is GGSGGGGSGGGGS. siRNA resistant APMAP (APMAP K347G) was made using site-directed mutagenesis.

All HyPer constructs (HyPer-Erlumen and HyPer-Cyto) and H_2_O_2_-insensitive HyPer mutant constructs (HyPer-Erlumen mutant and HyPer-Cyto mutant) were kindly provided to us by Prof. Geiszt (Semmelweis University, Hungary). Reticulon4a-eGFP was kindly provided by Gia Voeltz (Addgene 61807).^52^

For genetic manipulation of mouse APMAP in AML12 cells, we utilized CRISPR two based plasmid systems together with collaborator Dr. Joan Font-Burgada (FCCC). In one, we engineered a vector to express catalytically dead Cas9 (dCas9) fused with Dnmt3 and KRAB together with iCRE recombinase using the TBG hepatocyte-specific promoter. An additional cassette was included to express a gRNA targeting the mouse *APMAP* gene promoter. This enabled *APMAP* gene silencing as the dCas9-KRAB module targeted the APMAP gene promoter, inhibiting transcription (denoted as APMAP^KRAB^). Two other plasmids utilized Cas9-CRISPR and two different gDNAs targeted to *APMAP* to insert indels in the ORF (APMAP^KO_1^, APMAP^KO_2^). These were verified by Western blot to cause APMAP loss of expression.

### CRISPR gene editing

The APMAP C-terminal knock-in guide RNA sequence was designed using the online tool CRISPOR (Concordet, 2018) 5’-GCAGGGGCAGCTATCTGGGA-3’. The sequence was synthesized as oligos with BbsI overhangs and was cloned into the BbsI sites of pX459V2.0-HypaCas9^53^ (Addgene 108294). The homology repair template to make endogenous APMAP-mNeonGreen was made from two synthesized 800-bp homology regions gene fragments (Twist Biosciences) from upstream and downstream of the stop codon and the CDS of mNeonGreen cloned into pEGFP-N2 without its CMV promoter and GFP. The final sequence has a linker of GSFEFCSRRYRGPGIHRPVAT between the last amino acid of APMAP and the first amino acid of mNeonGreen.

U2OS cells were transfected with the pX459V2.0-HypaCas9 with the APMAP guide sequences inserted and with the APMAP-mNG repair template vector as above. After 24h, cells were treated with 3 ug/ml puromycin for 24h. Cells were grown up and then sorted for mNeonGreen fluorescence using a BD FACS Aria. Sorted cells were plated sparsely into a 10 cm culture plate and then grown until colonies reached >100 cells each. Colonies were picked individually using trypsin-coated sterile filter paper disks (Bel-Art F378470001) and plated in 24-well plates, where they were grown to confluence. Colonies were tested for homozygous APMAP knock-in using fluorescence imaging and Western blot.

### RNAi and transient transfections

Cells were plated in complete media and grown to a 50-60% confluency before conducting knockdown experiments in serum-free media. Cells were transfected with scrambled siRNA (Silencer Select negative control, Thermo Fisher 4390843 or Silencer Negative Control 1, Thermo Fisher AM4611) and APMAP siRNA (Silencer Select siRNA, Thermo Fisher s32757) with a final concentration of 10 nM, using Lipofectamine RNAiMAX reagent (Thermo Fisher 13778075) for 48 h. Cells were transfected with pre-incubated siRNA with a final concentration of 10 nM siRNA (3.75 μl of lipofectamine RNAiMAX in 125 μls of Opti-MEM, Gibco 31985-070) for 48 h. Cells were transfected again the following day with 10 nM siRNA for a 72 h total knockdown time. Media and reagents are scaled up and down based on surface area of plates. Tandem RNAi (10 nM)/plasmid transfection for rescue experiments were performed at 48 h mark using Lipofectamine 3000 (Invitrogen L3000015) as follows.

Plasmid transfection was performed using lipofectamine 3000 using 250 ng of DNA (split in half for dual transfection), 1 μl of P3000, 0.5 μl P3000, 0.75 μl Lipofectamine 3000 per 500 μl of total transfection/media mix. AML12 cells were transfected with APMAP^KO_1^, APMAP^KO_2^, and APMAP^KRAB^ plasmids using the above-mentioned method and incubated for 4 days to achieve effective gene knockout/ knockdown.

### ROS imaging and analysis

For BODIPY-C11^32,33^ imaging in U2OS cells, cells were transfected with control and APMAP siRNA as detailed above, with or without transfected APMAP-TagBFP as a co-transfection marker. Cells were treated with DMEM or DMEM with 250 μM tert-butyl hydroperoxide (TBHP) at 37°C for 2 h, and 10 μM BODIPY-C11 was added for the last 30 min of this treatment. Cells were then washed 3x with DMEM, fixed in 4% paraformaldehyde in PBS for 15 min, then washed 2x with PBS. Dishes were then stored at 4°C in the dark until imaged. BODIPY-C11 mammalian cell images were taken on a Nikon CSU-W1 spinning disk confocal microscope. For BODIPY-C11 quantification at the cell periphery, BFP-positive cells for each condition were chosen for analysis. 2-5 micron-length lines of 100-pixel width were drawn at the periphery of cells and quantified using the plot profile function in ImageJ. The local intensity maxima of the 510 nm-shifted fluorescence signal were reported.

For HyPer imaging in Huh7 cells, cells were transfected with HyPer-ERLumen, HyPer-ER-Lumen mutant, HyPer-Cytosolic, and HyPer-Cytosolic mutant, and imaged live on a Zeiss LSM880 laser scanning confocal equipped with Definite Focus for time lapse imaging. Pre-treatment images were taken, then cells were treated with 2 mM H_2_O_2_, and imaging was initiated after 10 s and continued at 5 s intervals for 6 min. HyPer-ER Lumen mutant imaging was used to visually verify that H_2_O_2_ treatment only changed the shift in fluorescence stimulation in the WT HyPer-ER-Lumen construct-expressing cells, not in those expressing the inactive mutant. For HyPer quantification, image background intensities were calculated using the average of 3 boxes in the cell-free area of the pre-treatment image. This background was subtracted from the rest of the images. A ROI was drawn manually around each cell, and integrated density and raw integrated density for 488- and 405-stimulated fluorescence were measured for each cell and time point. The ratio of 488-stimulated raw integrated density to the 405-stimulated raw integrated density was calculated for each time point/cell and plotted.

### Immunofluorescence staining

Cells were fixed with warmed 4% paraformaldehyde (PFA) solution in PBS for 10 min at room temperature (RT). For IF staining, fixed cells were rinsed with 1x PBS and permeabilized with 0.1% Triton X-100 in 1x PBS at RT for 5 min. Permeabilization was followed by blocking of cells in IF buffer (PBS containing 3% BSA and 0.1% Triton X-100) for 30 min. The cells were then incubated with primary antibody in IF buffer for 1 h and washed thrice with 1x PBS, followed by incubation with secondary antibody in IF buffer for 30 min. Post staining, cells were washed thrice with 1x PBS. Cells in PBS were either imaged immediately for microscopic studies or stored at 4°C. The primary antibodies used are mouse anti-HSP90B1 (1:100; Sigma-Aldrich; AMAb91019), rabbit anti-HSP90B1 (1:100; Sigma-Aldrich; HPA008424), rabbit anti-APMAP (1:100; Sigma-Aldrich; HPA012863), mouse anti-APMAP (1:150; Novus biologicals; NBP2-01716), rabbit anti-GFP (1:200; Abcam; ab290), rabbit anti-mCherry (1:200; Invitrogen; PA534974), rabbit anti-calnexin (1:500; Abcam; ab22595), mouse anti-CLIMP63 (1:100, Enzo, G1-296), Rabbit anti-calnexin (1:200; Abcam; ab22595), goat anti-Apolipoprotein B (1:200, Rockland immunochemicals, 600-101-111) and mouse anti-ceramide (1:50; Enzo; ALX-804-196). The secondary antibodies are donkey anti-mouse AF488 (Thermo Fisher Scientific; A21202), donkey anti-mouse AF594 (Thermo Fisher Scientific; A21203), donkey anti-rabbit AF488 (Thermo Fisher Scientific; A21206), donkey anti-rabbit AF594 (Thermo Fisher Scientific; A21207), donkey anti-goat AF488 (Thermo Fisher Scientific; A11055) and donkey anti-goat AF594 (Thermo Fisher Scientific; A11058) used at a dilution of 1:400 and incubated for 30 min. LDs were visualized by staining the cells with MDH (1:1,000; Abgent; SM1000a) for 15 min.

### Confocal fluorescence microscopy and Image analysis

Confocal microscopy of the cells was performed using the following equipment according to the experiment – a) 63x oil-immersion objective lens in a Zeiss laser scanning 780 confocal and Zeiss LSM880 Airyscan microscope; b) Live-cell imaging of endogenous APMAP using 4-100x objectives in Nikon TiE inverted microscope with a Yokogawa CSU-W1 spinning disk+SoRa module. All general image processing and final image preparation were done in ImageJ/Fiji. Statistical analysis for the data were done using GraphPad Prism 10.0.

#### Line scan analysis

After drawing an ROI onto the selected image with the line segment tool, line scan analysis was done for all channels separately using ‘plot profile’ function in ImageJ. The length and the pixel width of the line were kept consistent within an experiment. A high-resolution plot of the histogram displaying fluorescence intensity (Y-axis) of the signals along the distance of the line (microns) was exported for the appropriate markers used.

#### Quantification of LD area

LD area was measured using the Trainable WEKA Segmentation plugin from FIJI. Using max intensity projected images, boundaries of each cell were drawn using a free-hand tool. A WEKA classifier^54^ was trained to segment MDH-stained LDs from the cellular background. The modified images were then thresholded, which was followed by application of ‘analyse particles’ function. The average LD area covered per cell was then calculated in Excel. For the rescue experiments with overexpression constructs, transfected cells were selected using a freehand drawing tool, and then the area covered by LDs in each cell was calculated and plotted.

#### Quantification of ER morphology changes

Using ImageJ, images from each condition were converted to 8-bit maximum intensity projections. These 8-bit max projections were further subjected to unsharp masking with a radius of 2 and mask of 0.6. The max intensity projections were thresholded using the Otsu threshold^55^ to generate binary image. Each cell was manually traced using total ER fluorescent signal.

For ER morphological experiments, cells were stained with both α-calnexin and α-CLIMP-63. For percent occupancy of CLIMP63 within the total ER membrane (calnexin), the percent of both CLIMP63 and calnexin positive pixels per cell area were measured. For calculation of ER sheet expansion, ratio of % of CLIMP-63 positive pixels over % of calnexin positive pixels were measured. For measuring Anti-ceramide and anti-ApoB signals, % of ceramide-positive pixels or % of ApoB-positive pixels per cell were calculated.

### Electron microscopy

The U2OS cells were cultured under appropriate conditions on MatTek dishes and processed in the UT Southwestern Electron Microscopy Core Facility. The cells were fixed with 2.5% (v/v) glutaraldehyde in 0.1 M sodium cacodylate buffer, rinsed thrice in the buffer, and post-fixed in 1% osmium tetroxide and 0.8% K3[Fe(CN6)] in 0.1 M sodium cacodylate buffer at room temperature for 1 h. After rinsing the fixed cells with water, they were en bloc stained with 2% aqueous uranyl acetate overnight. On the following day, the stained cells were rinsed in buffer and dehydrated with increasing concentration of ethanol. Next, they were infiltrated with Embed-812 resin and polymerized overnight in a 60°C oven. A diamond knife (Diatome) was used to section the blocks on a Leica Ultracut UCT (7) ultramicrotome (Leica Microsystems), which were collected onto copper grids, and post stained with 2% aqueous uranyl acetate and lead citrate. A Tecnai G2 spirit transmission electron microscope (FEI) equipped with a LaB6 source and a voltage of 120 kV was used to acquire the images.

### RNA extraction and quantitative RT-PCR

Total RNA isolation from mammalian cells was performed according to the manufacturer’s instructions using Trizol reagent (Ambion Life Technologies; 15596026). RNA yields and purity were measured using a nanodrop. 2 μg of extracted RNA was used to generate cDNA using ReadyScript cDNA synthesis mix (Sigma; RDRT-100RXN) and the obtained cDNA were diluted 1:3 times with nuclease-free water for performing further experiments. qPCR was performed using SsoAdvanced Universal SYBR Green Supermix (BioRad; 172-5274) and the mRNA expressions were normalized to the housekeeping gene Cyclophilin A (CypA). All primers used for qPCR were Kicq pre-designed primers (Sigma).

### Fluorescence Protease Protection Assay

Followed previously established protocol.^56^ Briefly, 48 h after transient transfections with Reticulon4a-eGFP and APMAP^FL^-eGFP, Huh7 cells were washed with KHM buffer (110 mM potassium acetate, 20 mM HEPES, 2 mM MgCl_2_) at RT. The Labtek dish containing the cells were placed on microscope stage for fluorescence imaging and was imaged before the addition of digitonin to record the pre-permeabilisation stage. After determining 0.01% digitonin is sufficient for effective plasma membrane permeabilization, cells transfected with appropriate overexpression constructs were then treated with KHM buffer containing 0.01% digitonin for 1 minute followed by KHM buffer containing 50 μg/ml Proteinase K. Images of these cells from 30 seconds to 8 minutes or until complete loss of fluorescence signal were recorded using live cell imaging. Each condition was imaged in triplicates.

### Thin Layer Chromatography

A protocol adapted from Bligh and Dyer, 1959^57^ was used to extract lipids from cultured cells. Briefly, cells cultured in complete media were washed twice in PBS, scraped, and collected, and their weights were measured. The cell pellets were lysed by vortexing in the presence of chloroform, followed by methanol and 500 mM NaCl in 0.5% acetic acid to the final concentration such that the ratio of chloroform: methanol: water was 2:1:0.8. The suspension was centrifuged at 4,000 rpm for 15 min at 4°C and the bottom chloroform layer, comprising the lipids, was collected. This lipid layer was dried and then again resuspended in chloroform to a final concentration normalized to the initial cell pellet weight. Extracted lipids alongside serially diluted standard neutral lipids of known concentrations were separated on TLC plates using hexane: diethyl ether: acetic acid solvent (80:20:1, v/v/v). TLC plate was air dried for 10 min, spray stained with 3% copper (II) acetate in 8% phosphoric acid and incubated at 145°C in the oven for 1 h to allow bands to develop for scanning and imaging. Neutral lipid (TAG) band intensity was quantified using Fiji/ImageJ software, and lipid concentrations were calculated from the standard curve generated with standard mixture.

### Proteomic analysis

To prepare samples of control and APMAP-TurboID-mNeonGreen overexpressing cells for biotin-enrichment and proteomic analysis^58^, Huh7 cells were transfected with APMAP-TurboID-mNeonGreen for 24 h. Transfected and untransfected cells were treated with DMEM +/- 50 μM biotin for 10 min, then put on ice and washed with cold PBS 2x. Cells were collected by scraping and lysed with RIPA buffer with Halt protease inhibitor cocktail (Thermo; 78429) for 15 min on ice and pushed through a 25G needle 5x. The sample was centrifuged at 13000xg for 10 min at 4°C; protein in the supernatant was quantified using the Pierce Bradford assay. 10% of samples were saved for western blot of the input, and 90% of samples were dialyzed using 3.5 MWCO Slide-A-Lyzer cassettes (Thermo Scientific; 66330) in RIPA buffer with gentle stirring overnight at 4°C. Protease inhibitor was then added to the sample, and protein was re-quantified. Beads were incubated with RIPA-washed streptavidin magnetic beads (Thermo Fisher 88817) (200 μl beads per 1.57 mg protein) with 500 µl additional RIPA buffer at 4°C overnight with rotation. Beads were pelleted with a magnetic rack (with the flow-through saved), washed with detergent-free RIPA for 2 min, then washed with 1 M KCl for 2 min, then washed with 0.1 M Na_2_CO_3_ for 10 s, then washed with 2 M urea in 10 mM Tris-HCl pH 8 for 10 s, then washed 2x with detergent-free RIPA buffer, then suspended in detergent-free RIPA buffer and delivered to the UT Southwestern Proteomics Core Facility. The buffer was removed from the vial containing the streptavidin beads, and the beads were washed with PBS. To the beads, 100 μl of 2M urea and 100 mM Tris-HCl was added. Samples were reduced with 4ul of 500mM tris(2-carboxyethyl) phosphine (TCEP) for 30 min at room temperature. The free cysteines were then alkylated by adding 4 μl of 500 mM iodoacetamide (IAA) and incubated in the dark at room temperature for 30 min. The buffer was removed and 100 μl of Tris-HCl was added to the beads along with 1 μg of trypsin. The samples were digested overnight at 37°C with shaking.

Following digestion, the supernatant from each sample containing the peptides was transferred to a new vial. 100 μl of 0.1% TFA was added to the vials containing the beads, shaken for 5 min, and the solution was transferred to the corresponding vials containing the supernatants. These samples were then cleaned using an Oasis HLB microelution plate (Waters), and dried in a SpeedVac. 51 μl of TEAB was added to each sample, and 20 μl of TMT 10plex reagent (Thermo) was added to each tube (Reagents TMT-130C and 131 not used). Samples were incubated for 1 h for the TMT labelling, and the reaction was quenched with 5% hydroxylamine for 20 minutes. The TMT-labeled samples were combined and dried in a SpeedVac. The peptides were reconstituted in enough volume of 2% acetonitrile, 0.1% TFA to get a concentration of ∼0.75 μg/μl by NanoDrop A205. 2 μl of this sample was injected onto an Orbitrap Fusion Lumos mass spectrometer coupled to an Ultimate 3000 RSLC-Nano liquid chromatography system. Samples were injected onto a 75 μm i.e., 75-cm long EasySpray column (Thermo) and eluted with a gradient from 1-28% buffer B over 180 min, followed by 28-45% buffer B over 25 min. Buffer A contained 2% (v/v) ACN and 0.1% formic acid in water, and buffer B contained 80% (v/v) ACN, 10% (v/v) trifluoroethanol, and 0.1% formic acid in water. The mass spectrometer operated in positive ion mode with a source voltage of 2.0 kV and an ion transfer tube temperature of 300°C. MS scans were acquired at 120,000 resolution in the Orbitrap over a mass range of m/z = 400-1600, and top speed mode was used for SPS-MS3 analysis with a cycle time of 2.5 s. MS2 was performed using collisionally-induced dissociation (CID) with a collision energy of 35% for ions with charges 2-6. Dynamic exclusion was set for 25 s after an ion was selected for fragmentation. Real-time search was performed using the reviewed human protein database from UniProt, with carbamidomethylation of Cys and TMT 6plex modification of Lys and peptide N-termini set as static modifications and oxidation of Met set as a variable modification. Two missed cleavages and up to 3 modifications per peptide were allowed. The top 10 fragments for MS/MS spectra corresponding to peptides identified by real-time search were selected for MS3 fragmentation using high-energy collisional dissociation (HCD), with a collision energy of 58%.

Raw MS data files were analyzed using Proteome Discoverer v2.4 SP1 (Thermo), with peptide identification performed using Sequest HT searching against the human protein database from UniProt. Fragment and precursor tolerances of 10 ppm and 0.6 Da were specified, and three missed cleavages were allowed. Carbamidomethylation of Cys and TMT 10plex labelling of N-termini and Lys sidechains were set as fixed modifications, with oxidation of Met set as a variable modification. The false-discovery rate (FDR) cutoff was 1% for all peptides.

### ELISA for determining ApoB levels in culture media

Previously established sandwich ELISA protocol for quantification of ApoB secretion was followed.^29^ Briefly, after 72 h of siRNA transfections, control and APMAP-depleted Huh7 cells were treated with OA for 16 h. Following this, the cells were washed, topped with fresh serum-free media and subsequently both media and cells were collected at 0 h, 2 h, 4 h, 8 h and 24 h. Rabbit polyclonal ApoB antibody (1:200; Abcam) was used as the capture antibody and goat polyclonal ApoB antibody (1:200, Rockland) was used as detection antibody. Horse radish peroxidase (HRP)-conjugated anti-goat IgG secondary antibody (1:500; Thermo Fisher; PA128664) was used to detect the capture antibody-antigen complex. To detect HRP, 100 μl of tetramethylbenzidine (TMB) peroxidase substrate (Millipore Sigma; T0440) was added and allowed to react for 4 mins for color development. The color reaction was stopped by addition of 100 μl 1M HCl, and the absorbance was read at 450 nm using a Molecular Devices VersaMax absorbance microplate reader. Standard curves were prepared using purified human VLDL (Millipore Sigma, LP1) diluted in same cell culture media and were used for determining the concentration of ApoB in the samples.

### Targeting Zebrafish ***apmap*** with CRISPR/Cas9

Transgenic LipoGlo ApoBb.1-NanoLuc zebrafish larvae (ApoBb.1Nluc/Nluc) were used for CRISPR injections at the one cell stage. Each larvae was injected with four guide RNA’s all targeting the same gene^59^ or four guides that have a scrambled sequence that is not predicted to target a gene or have previously exhibited any phenotype. As an additional control for viability, a subset of larvae from each clutch was set aside and not injected.

### Zebrafish LipoGlo Assays

Zebrafish larvae were imaged via chemiluminescence microscopy at 3 and 5 days post fertilization (dpf) as described.^15^ Total larval ApoB levels were determined from tissue extracts derived by sonication of larvae and their total LipoGlo-NanoLuc luminescence quantified. Specifically, ApoBb.1-NanoLuc levels were assayed as described^15^ with one modification of using 4uL of tissue homogenate mixed with 76uL x2 NanoLuc buffer in a 96-well black Perkin-Elmer OptiPlate. The quantification of lipoprotein size distribution and all whole-mount chemiluminescent imaging was done according the Thierer et al.^15^

### Lipidomics

The solvents used were either HPLC or LC/MS grade and purchased from Sigma-Aldrich (St Louis, MO, USA). Splash Lipidomix stadndards were purchased from Avanti (Alabaster, AL, USA). All lipid extractions were performed in 16×100mm glass tubes with PTFE-lined caps (Fisher Scientific, Pittsburg, PA, USA). Glass Pasteur pipettes, eVol automated analytical syringes (Trajan, Australia) and solvent-resistant plasticware pipette tips (Mettler-Toledo, Columbus, OH, USA) were used to minimize leaching of polymers and plasticizers.

The lipid liquid-liquid extraction was performed using a modified methyl-tert-butyl ether (mTBE) method.^60^ Briefly, an aliquot containing 250K cells was transferred to a glass tube where 1 ml of water, 1 ml of methanol and 2 ml of mTBE were added. The mixture was vortexed, then centrifuged at 2671*×g* for 5 min, the organic phase was collected, spiked with 20 μl of a 1:5 diluted Splash Lipidomix standard solution and dried under N_2_ air flow. The samples were resuspended in hexane for lipidomics analysis.

The lipid profile was obtained by LC-MS/MS using a SCIEX QTRAP 6500^+^ (SCIEX, Framingham, MA) equipped with a Shimadzu LC-30AD (Shimadzu, Columbia, MD) high-performance liquid chromatography (HPLC) system and a 150×2.1 mm, 5 μm Supelco Ascentis silica column (Supelco, Bellefonte, PA). Samples were injected at a flow rate of 0.3 ml/min at 2.5% solvent B (methyl tert-butyl ether) and 97.5% Solvent A (hexane). Solvent B was increased to 5% over 3 min and then to 60% over 6 min. Solvent B was decreased to 0% during 30 sec while Solvent C (90:10 (v/v) isopropanol-water) was set at 20% and increased to 40% during the following 11 min. Solvent C is increased to 44% over 6 min and then to 60% over 50 sec. The system was held at 60% solvent C for 1 min prior to re-equilibration at 2.5% of solvent B for 5 min at a 1.2 ml/min flow rate. Solvent D [95:5 (v/v) acetonitrile-water with 10 mM Ammonium acetate] was infused post-column at 0.03 ml/min. Column oven temperature was 25°C. Data was acquired in positive and negative ionization mode using multiple reaction monitoring (MRM). The LC-MS/MS data was analyzed using MultiQuant software (SCIEX). The identified lipid species were normalized to its corresponding internal standard.

### Fly Cultures and Strains

*Drosophila* L3 late feeding larvae and 7-10 day old adults were used for larval and adult assays, respectively. All *Drosophila* animal research was performed in accordance with NIH recommended policies. Flies were reared and maintained on standard fly food containing cornmeal, yeast, molasses (∼5% sugars), and agar. The *Dcg-Gal4* fat body (FB)-specific driver, previously described (Suh et al., 2008) was provided by Jonathan M. Graff (UTSW). The *Cg-Gal4;UAS-Dcr2* FB driver was received from Steve Jean (Université de Sherbrooke). Ubiquitous driver *Da-Gal4*, TRiP *UAS-RNAi* stocks for Hmu (#41627, #41915), and ER organelle marker *UAS-Sec61B-TdTomato* (#64747) used were obtained from the Bloomington Stock Center (Bloomington, IN). All Gal4/UAS crosses, including those targeting Hmu knockdown were cultured at 25°C to maintain uniform transgene expression. Crosses involving *Cg-Gal4;UAS-Dcr2* with RNAi lines were cultured at 29°C to maximize knockdown efficiency.

### Fat Body/Larval Gut Dissection and LD Staining

Late feeding L3 larvae were gently removed from inside the food media using a paintbrush and rinsed in water to remove food particles. Larval fat bodies (FBs) or guts were dissected in PBS using Dumont #5 forceps (Electron Microscopy Sciences). All tissues were fixed in 4% paraformaldehyde for 20 min at room temperature and rinsed briefly in PBS prior to staining. Lipid droplets in FBs were stained by incubating FBs in 100 μM monodansylpentane (MDH) LD stain (Abgent) for 30 min at RT, in the dark. LDs in guts were stained with 1 µM Nile Red for 30 min at RT, in the dark. Next, FBs and guts were rinsed in PBS and mounted on slides in SlowFade Gold antifade reagent with DAPI (Invitrogen) for subsequent imaging.

### BODIPY-C11 lipid peroxidation

Fat bodies dissected from late feeding larvae were fixed in 4% paraformaldehyde for 20 min, rinsed briefly in PBS, then incubated in 2 μM BODIPY-C11 581/591 (Invitrogen, D3861) in the dark at room temperature for 30 min. After rising with PBS, fat bodies were mounted on slides in SlowFade Gold antifade reagent with DAPI (Invitrogen). Tissues were imaged by confocal fluorescence microscopy (40X oil) using both RFP and GFP filters to detect non-oxidized and oxidized state. The oxidized/reduced ratio was quantified.

### Confocal Fluorescence Microscopy

Prepared slides were imaged using Zeiss LSM880 inverted laser scanning confocal microscope. Tissues stained with organelle dyes or expressing transgenic fluorescent-tagged proteins (e.g., *UAS-Hmu-GFP*, *UAS-Sec61B-TdTomato*) were imaged using the appropriate channel filter DAPI/FP/GFP. FBs (MDH) were imaged using a 40X oil immersion objective, as 1 um z-stack sections. Larval guts (Nile Red) were imaged using a 20X objective.

### Thin Layer Chromatography (TLC)

Larvae were weighed, then homogenized in 2:1:0.8 of methanol:chloroform:water. Samples were incubated in a 37°C water bath for 1 h. Chloroform and 1 M KCl (1:1) were added to the sample, centrifuged at 3000 rpm for 2 min, and the bottom layer containing lipids was aspirated using a syringe. Lipids were dried using argon gas and resuspended in chloroform (100 μl of chloroform/7mg of fly weight). Extracted lipids alongside serially diluted standard neutral lipids of known concentrations were separated on TLC plates as explained above. In the case of dissected larval guts, dried lipids for all samples were re-suspended in 120 μl of chloroform and final lipid concentrations were calculated by normalizing to protein in sample, measured by standard Bradford assay. Hemolymph (4 μl/sample) for DAG measurements collected from larvae was diluted in 0.1% N-Phenylthiourea (Sigma-Aldrich) in 50 μl PBS. The same lipid extraction protocol was followed, dried lipids were re-suspended in 75 µl of chloroform, and equal volumes were loaded on TLC plates.

### *Drosophila* Tissue Immunofluorescence

FBs dissected from larvae were fixed in 4% paraformaldehyde at room temperature for 30 min on a nutator, then washed 4 times with 1 X PBST (PBS in 0.1% Tween), 10 min each. Tissues were blocked in 1 X PBST + 2% normal goat serum for 2 hours at room temperature, then incubated with Rb-ApoII primary antibody (1:500 dilution, provided by Dr. Akhila Rajan) added to the blocking buffer, overnight at 4°C. Next, FBs were washed 4 times with 1 X PBST 15 min each, followed by incubation with anti-rabbit Alexa546 secondary antibody (1:1000 dilution) in blocking buffer at room temperature in the dark for 2 h on a nutator. FBs were rinsed again 4 times in 1 X PBST 15 min each, and subsequently mounted in SlowFade Gold antifade reagent with DAPI. Fluorescent structures were observed and imaged by confocal microscopy using standard RFP channel filter at 1 μm step-sizes.

### Statistical Analysis

All experiments were conducted at least three independent times to provide scientific rigor. Appropriate sample sizes and controls, including negative control when appropriate were applied for all microscopic and biochemical experiments. Student t-tests or ANOVA tests were used for comparative statistics or multiple variable experiments respectively. Statistical significances were applied for P<0.05.

## Supplementary figures

**Figure S1.**
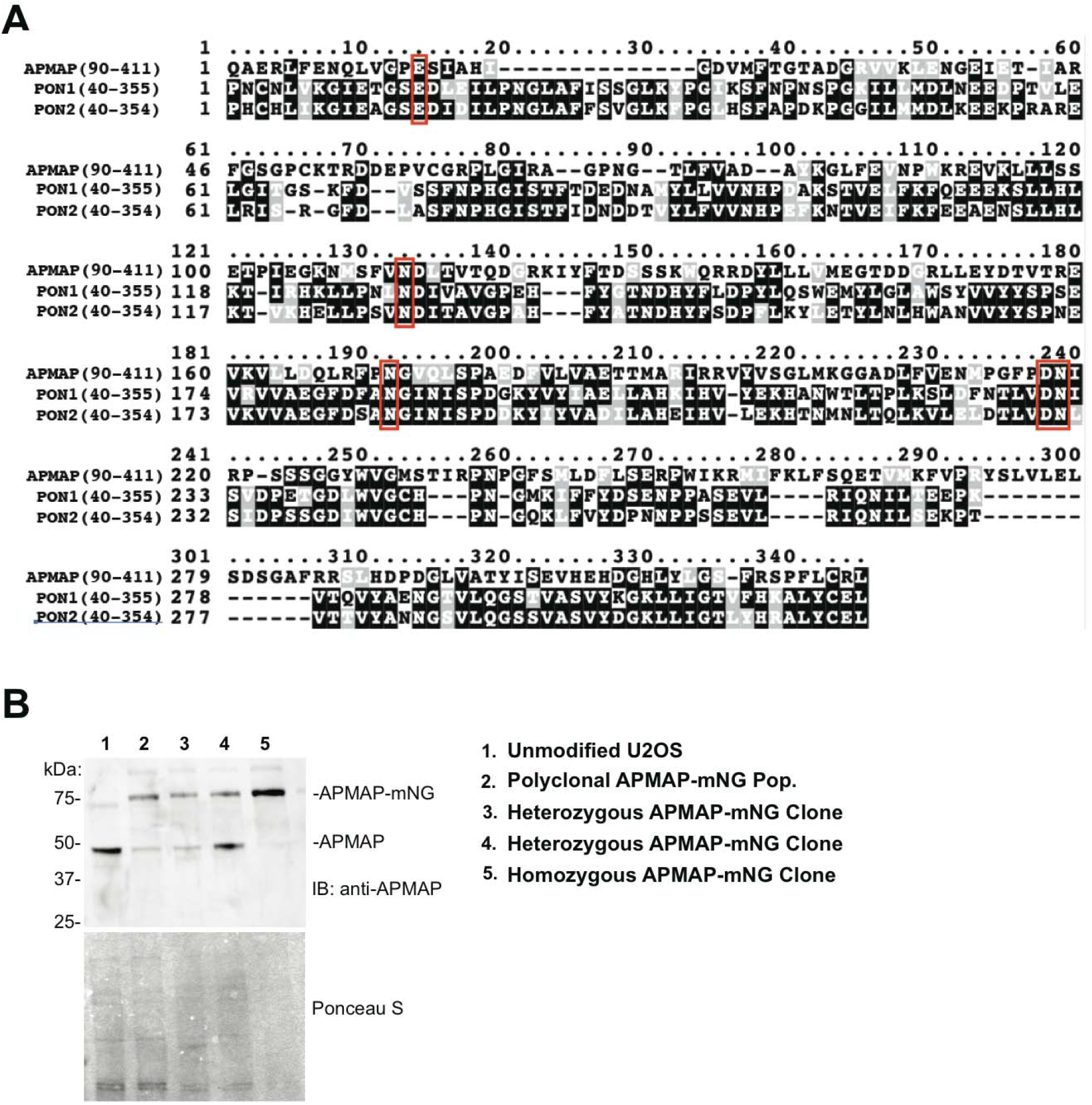
A. Amino acid alignment of 6-bladed b-propeller regions of APMAP with PON1 and PON2. Black background represents perfectly conserved amino acids. Grayed background represents conservation in similar amino acids. Alignment and color coding done using T-COFFEE multiple sequence alignment program and Boxshade respectively B. Validation of CRISPR-Cas9 integrated homozygous mNeonGreen knock-in APMAP clonal U2OS cell lines using western blot

**Figure S2.**
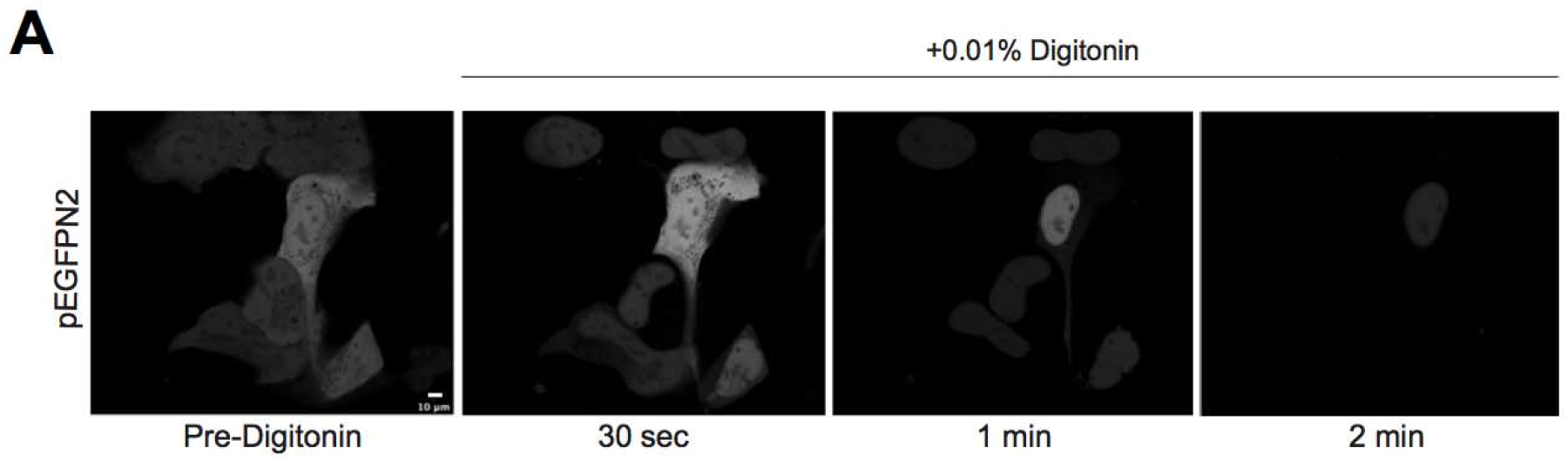
A. Live cell confocal micrographs of U2OS cells transfected with pEFGP-N2 expressing only EGFP, treated with 0.01% digitonin to check effective permeabilization of plasma membrane. Images taken before and after addition of digitonin. Scale bar = 10 um

**Figure S3.**
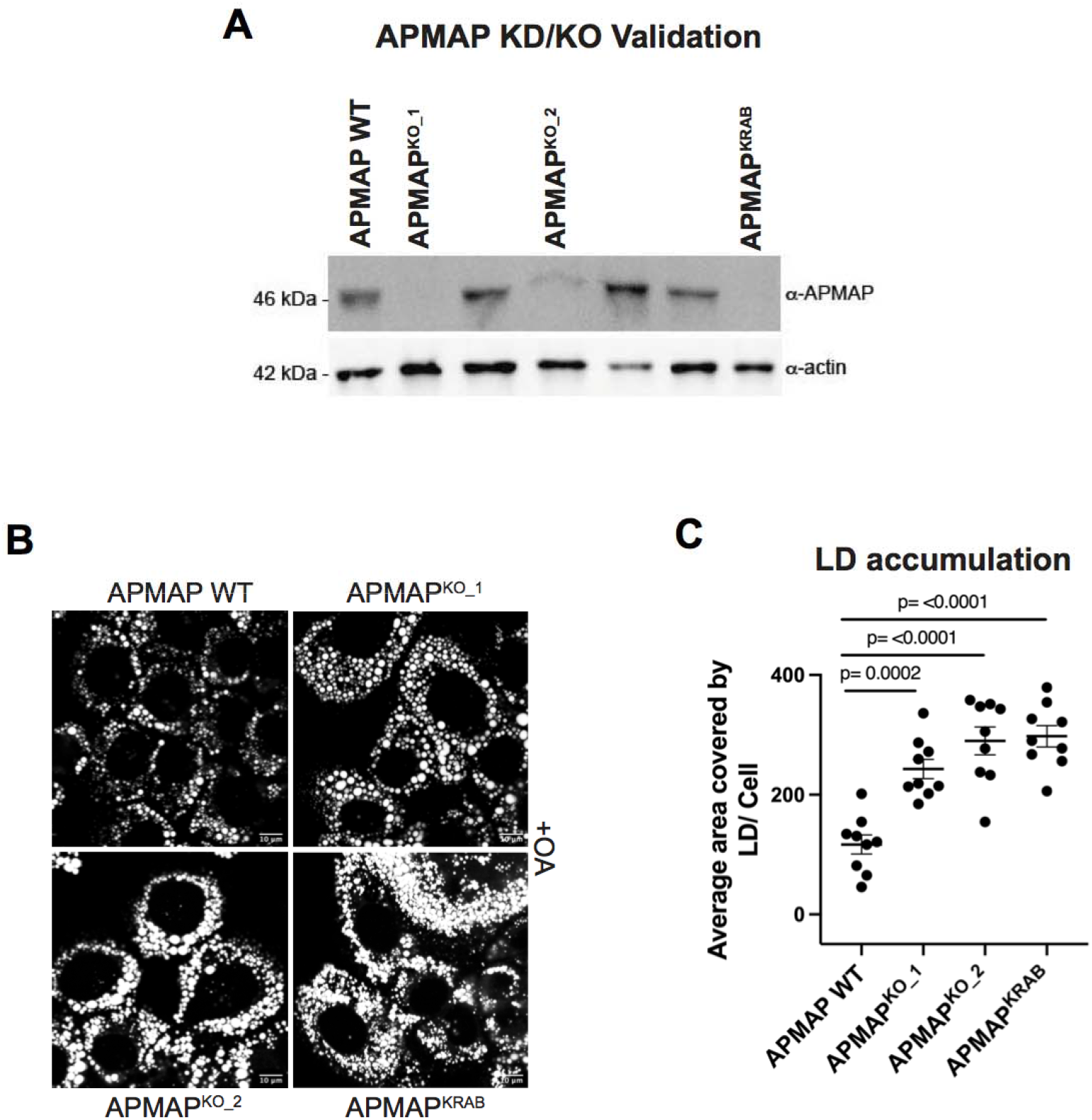
A. Western blot validation of efficient APMAP knockout/ knockdown in AML12 cells using plasmid transfections of KO (Cre) and KRAB constructs after 96 h incubation B. Confocal micrographs showing LD accumulation in WT, APMAPKO and APMAPKRAB transfected AML12 cells. LDs were visualized by MDH (gray). Scale bar= 10 μm C. Scatter dot plot representing quantified average area covered by LDs per cell. Total LD area was derived from 9 fields of view, each consisting of 10-12 cells and from three sets of experiments (Ordinary one-way ANOVA with Sidak’s multiple comparisons; α = 0.05). Means +/- SEMs shown

**Figure S4.**
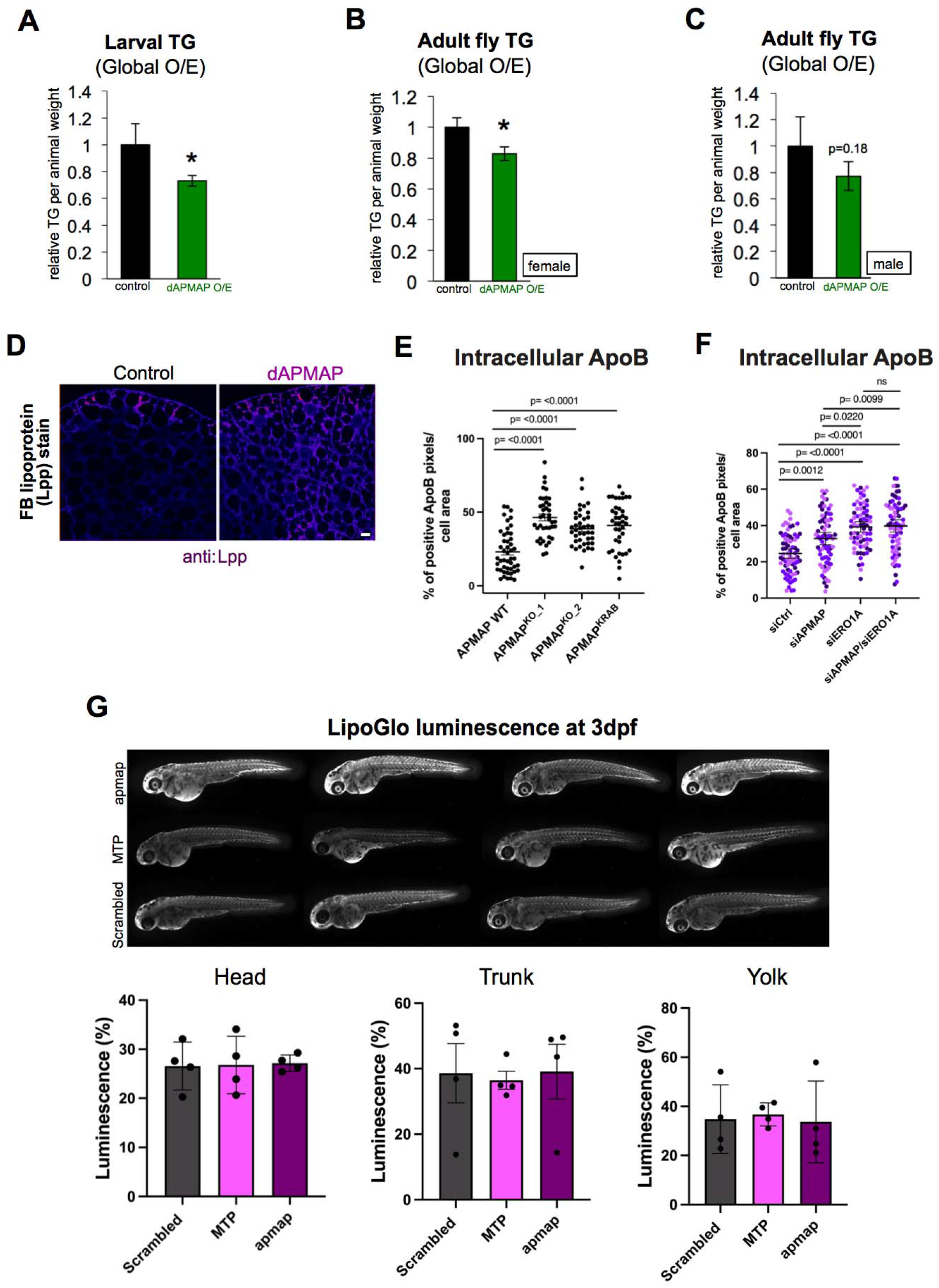
A. TLC quantification of relative female whole-larval gut triglyceride (TG) levels from control (*Da-Gal4*) and dAPMAP-eGFP global overexpression. N=3; *P<0.0322; Two tailed unpaired t-test B. TLC quantification of female flies with relative triglyceride (TG) levels from control (*Da-Gal4*) and dAPMAP-eGFP global overexpression. N=3; p=0.18; Two tailed unpaired t-test C. TLC quantification of male flies with relative triglyceride (TG) levels from control (*Da-Gal4*) and dAPMAP-eGFP global overexpression. N=3; p=0.18; Two tailed unpaired t-test D. Confocal micrographs of fat body tissue sections from control (*Dcg-Gal4*) and dAPMAP RNAi line with FB-specific RNAi depletion. Tissue was stained with α- LPP (lipophorin) demonstrating lipoprotein levels in each condition. Scale bar = 10 mm E. Scatter dot plot depicting intracellular ApoB levels for wildtype (WT) AML12 cells and AML12 cells with CRISPR-Cas9 mediated KO of APMAP (KO_1, KO_2) as well as dCas9-KRAB mediated silencing of APMAP (APMAP^KRAB^). Each dot represents the percent area of ApoB positive pixels per cell. (n=40; N=3; Ordinary one-way ANOVA with Sidak’s multiple comparisons; α = 0.05; Means +/- SEMs shown) F. Scatter dot plot of intracellular ApoB levels in Huh7 cells treated with siCtrl, siAPMAP, siERO1A and siAPMAP/siERO1A treated different conditions. Each dot depicts the percent area of ApoB positive pixels per cell. (n=70; N=3; Ordinary one-way ANOVA with Sidak’s multiple comparisons; α = 0.05) G. LipoGlo luminescence micrographs of 3 days post fertilization (3dpf) showing distribution of ApoB-containing lipoproteins (LipoGlo) in zebrafish larval vasculature. Animals were either treated with non-targeting CRISPR-based guides (scrambled) or gRNAs directed to delete zebrafish APMAP or MTP. Luminescence quantifications for head, trunk, and yolk displayed from N=4 independent experiments.

**Figure S6.**
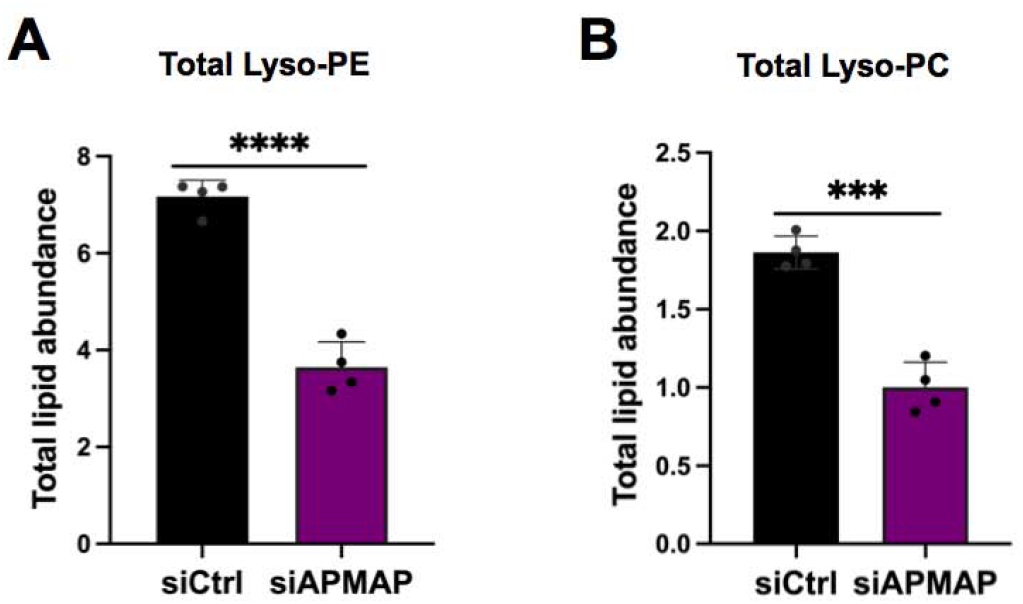
A. Total lipid abundance of Lyso-PE in siCtrl and siAPMAP treated Huh7 cells, measured with LC-MS/MS. N=4; ****P<0.0001, Two-tailed unpaired t-tests B. Total lipid abundance of Lyso-PC in siCtrl and siAPMAP treated Huh7 cells, measured with LC-MS/MS. N=4; ***P<0.0002, Two-tailed unpaired t-tests

**Figure S7.**
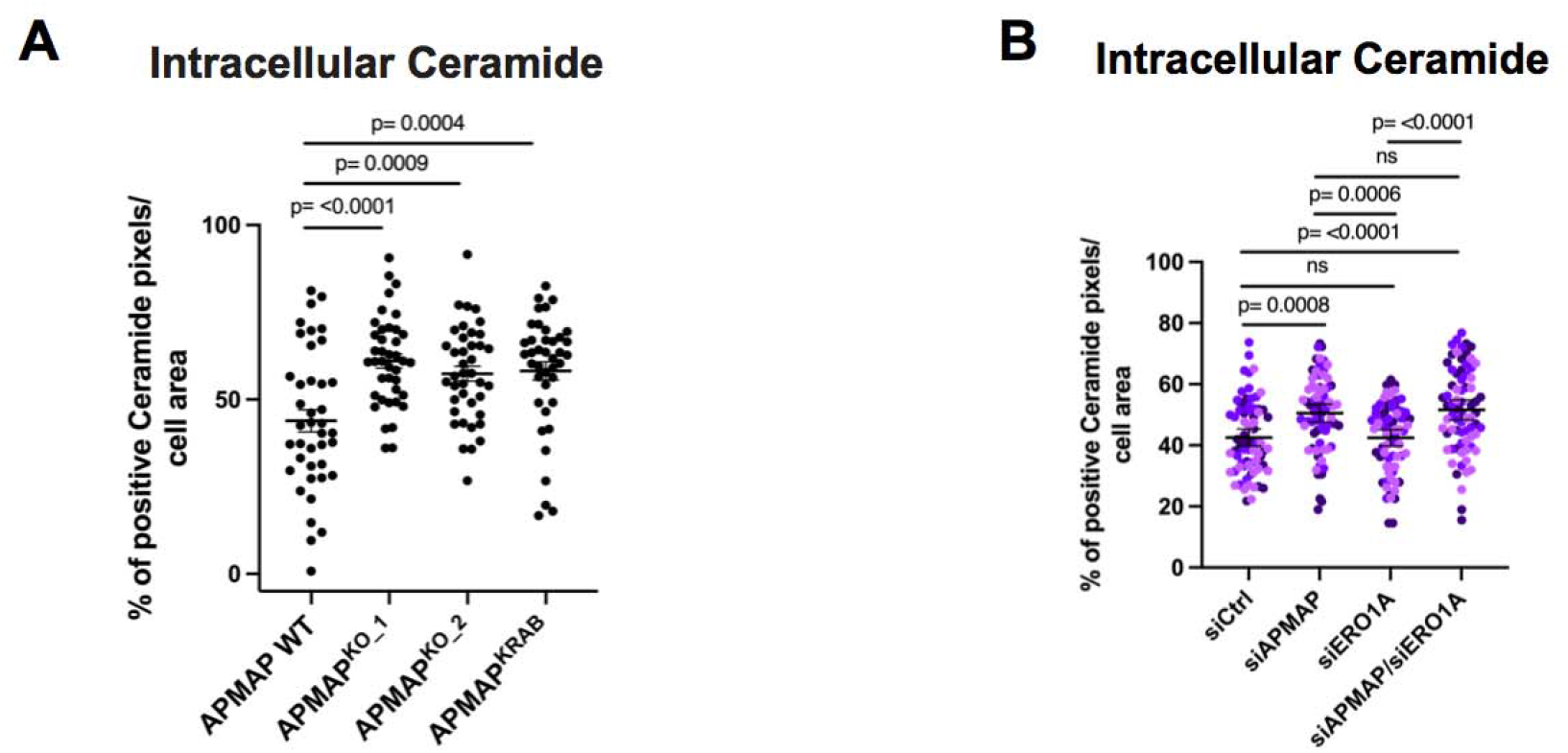
A. Scatter dot plot depicting intracellular ceramide quantifications in either wildtype (WT) AML12 cells or AML12 cells with CRISPR-Cas9 mediated KO of APMAP (KO_1, KO_2) as well as dCas9-KRAB mediated silencing of APMAP (APMAP^KRAB^). Each dot represents the percent area of ceramide positive pixels per cell. (n=40; N=3; Ordinary one-way ANOVA with Sidak’s multiple comparisons; α = 0.05; Means +/- SEMs shown) B. Scatter dot plot of intracellular ceramide levels in Huh7 cells treated with siCtrl, siAPMAP, siERO1A and siAPMAP/siERO1A treated different conditions. Each dot depicts the percent area of ceramide positive pixels per cell. (n=73; N=3; Ordinary one-way ANOVA with Sidak’s multiple comparisons; α = 0.05)

